# Learning-Dependent Shift from Right to Left CA3 Input Dominance Shapes the Evolution of Right CA1 Spatial Maps

**DOI:** 10.1101/2025.10.26.684700

**Authors:** Anqi Jiang, Douglas GoodSmith, Julliana Ramirez-Matias, Ariana F. Tortolani, Mark E. J. Sheffield

## Abstract

CA1 place fields support spatial maps critical for memory, yet how bilateral CA3 inputs shape these maps during learning remains unclear. Using two-photon calcium imaging and optogenetic inhibition in head-fixed mice navigating a virtual track, we examined left and right CA3 projections to right CA1 (CA1_R_) as animals familiarized to a novel environment. CA1_R_ maps were initially inaccurate but stabilized after ∼10 laps, defining an early-phase of map refinement followed by a late-phase of stability. During the early-phase, right CA3 inputs predominantly drove refinement, whereas left CA3 inputs controlled stability later. These effects arose at the single-cell level, with right CA3 inputs driving high-amplitude, reliable fields early and left inputs supporting reliable fields later. Axonal recordings revealed a matching shift: right CA3 axons showed greater place-field activity and reliability early, whereas left CA3 axons became more reliable later. Thus, CA3 input dominance transitions from right to left, coordinating CA1_R_ map refinement and stabilization.

## Introduction

The hippocampus is essential for spatial and episodic memory, with dorsal CA1 place cells forming the neural basis of a spatial representation ^1,2^. These neurons fire at specific locations within an environment, collectively generating a cognitive map of space ^3^. Understanding how these representations emerge and evolve over time is critical for uncovering the neural mechanisms underlying memory formation, retrieval, and updating.

Familiarization to a novel environment is a form of incidental spatial learning that occurs within a few minutes of exploration ^4–6^. During this period, CA1 exhibits an increase in place field formation ^7^, and at the population level, neural activity gradually evolves into a spatially structured state that is relatively stable ^8–11^. Associated with this period, dendritic-targeting interneurons in CA1 show reduced activity that enhances dendritic spike prevalence that can boost synaptic potentiation ^12,13^. Over the next few minutes and trials as animals become more familiar with their environment, interneuron activity gradually increases, dendritic spike frequency declines, and inhibitory circuits develop inverse spatial tuning relative to place cells ^12,13^. Moreover, behavioral timescale synaptic plasticity (BTSP) ^14,15^—facilitating rapid place field formation and shifting of place fields— is highest during the initial moments in a novel environment but gradually diminishes with familiarity ^16,17^. Together, these observations support two distinct phases in CA1 during exploration of a novel environment: an early phase of spatial representation formation and refinement, followed by a late phase of stabilized representations. But how inputs into CA1 contribute to these early and late phases is not clear.

Bilateral CA3 provides the primary excitatory input to CA1 and is essential for spatial learning ^18–20^. Left and right CA3 inputs form synapses that exhibit molecular and physiological differences that may influence their functional contributions ^21–29^. Synapses from Left CA3 inputs onto bilateral CA1 pyramidal cells contain more GluN2B-enriched NMDARs, which facilitate synaptic potentiation, whereas right CA3 synapses have a higher density of AMPARs, supporting stronger baseline transmission ^22,28^. Electrophysiological studies suggest that long-term potentiation (LTP) is more readily induced at left CA3-CA1 synapses under specific conditions, while right CA3-CA1 synapses do not undergo LTP under those same conditions ^21,23^. Behaviorally, inhibiting left CA3 disrupts long-term spatial memory, whereas right CA3 inhibition has minimal effects ^23,25^. These findings suggest that left and right CA3 inputs may differentially contribute to CA1 spatial dynamics across distinct memory phases.

To address this, we used two-photon calcium imaging to record right CA1 (CA1_R_) place cell activity while selectively inhibiting left or right CA3 inputs at specific timepoints during and following learning: during initial novel exploration (first 3 mins) when behavioral measures show ongoing learning and CA1 representations are forming and refining; after 3 mins when behavior shows no further learning and CA1 representations have stabilized (Fig. 1, 2). Additionally, we recorded CA3 axon activity in CA1 to examine how left and right CA3 inputs evolve over time during the early and later phases, providing insight into how they influence CA1_R_ dynamics.

**Fig 1.**
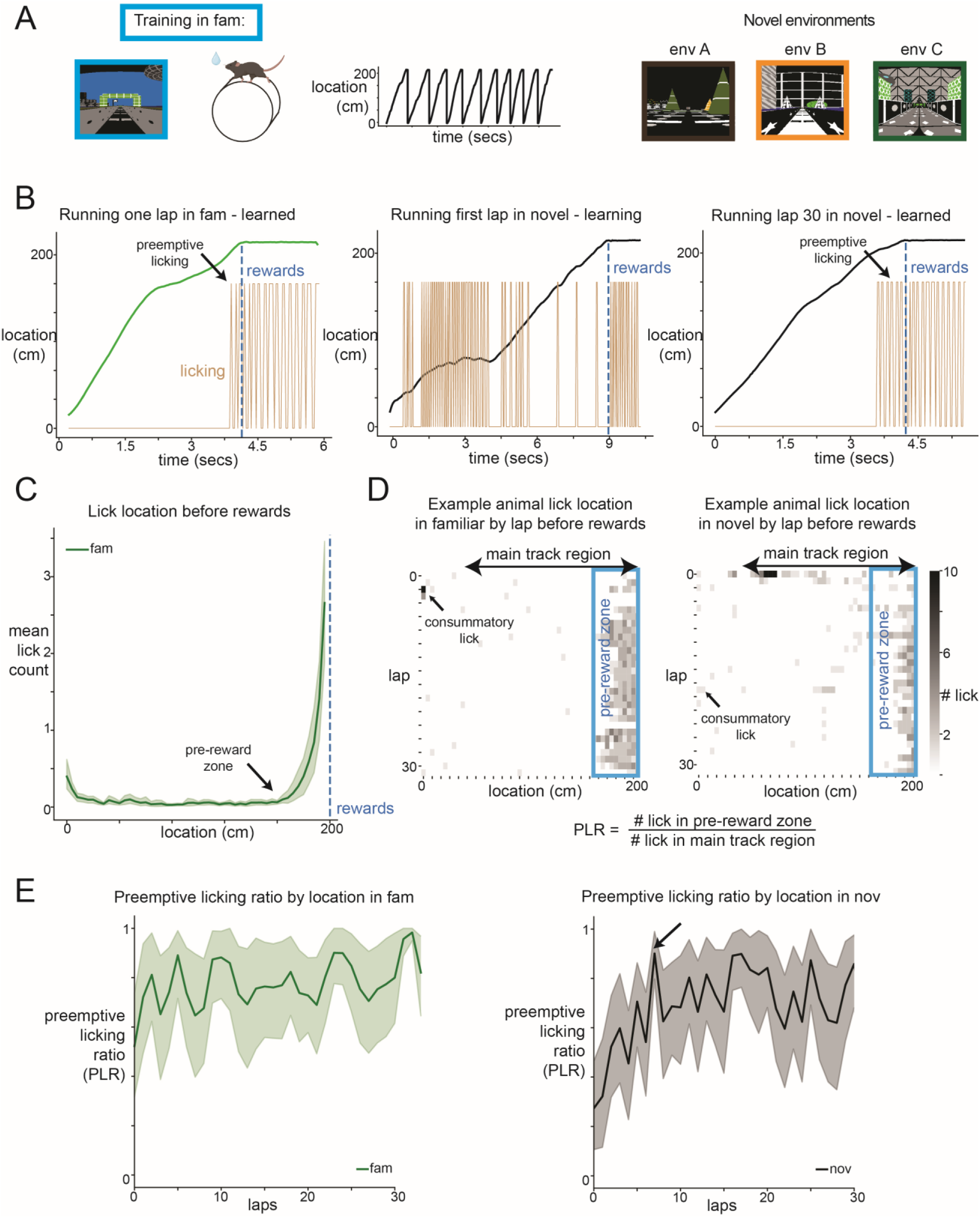
Anticipatory licking as a behavioral readout of spatial learning and familiarization to a novel environment. A. Behavioral training setup: mouse running on the treadmill in a 2m track VR environment. left: familiar VR environment. right: 3 different novel VR environments. B. Example animal running one lap in left: familiar env (learned), middle: first lap in novel env (during learning), and right: lap 30 in novel env (learned). Over repeated laps, animals develop preemptive licking anticipating water rewards prior to reaching the rewards zone. C. Mean lick count by location on the track before reward delivery from all animals (n=12). The pre-reward zone (start of zone indicated by arrow) was defined as the start of elevated lick ratio in the familiar environment identified by the kneedle algorithm (see Methods, Supp Fig8). Consummatory lick count after reward delivery was removed. D. Example animal lick location on the track by lap in familiar (left) and novel (right) environments. Consummatory licks often happen at the beginning of the track as animals finish consuming water rewards from the previous lap. We defined the “main track region” to include track locations after consummatory licks and before reward delivery. Preemptive licking ratio (PLR) is defined as the ratio of number of licks in pre-reward zone over number of licks in the main track region. E. PLR over laps in familiar (left) and novel (right) environments averaged across all animals (n=12). Arrow points at the transition lap (lap 8) from continuous rising to stability in the novel environment identified by the kneedle algorithm.

**Fig 2.**
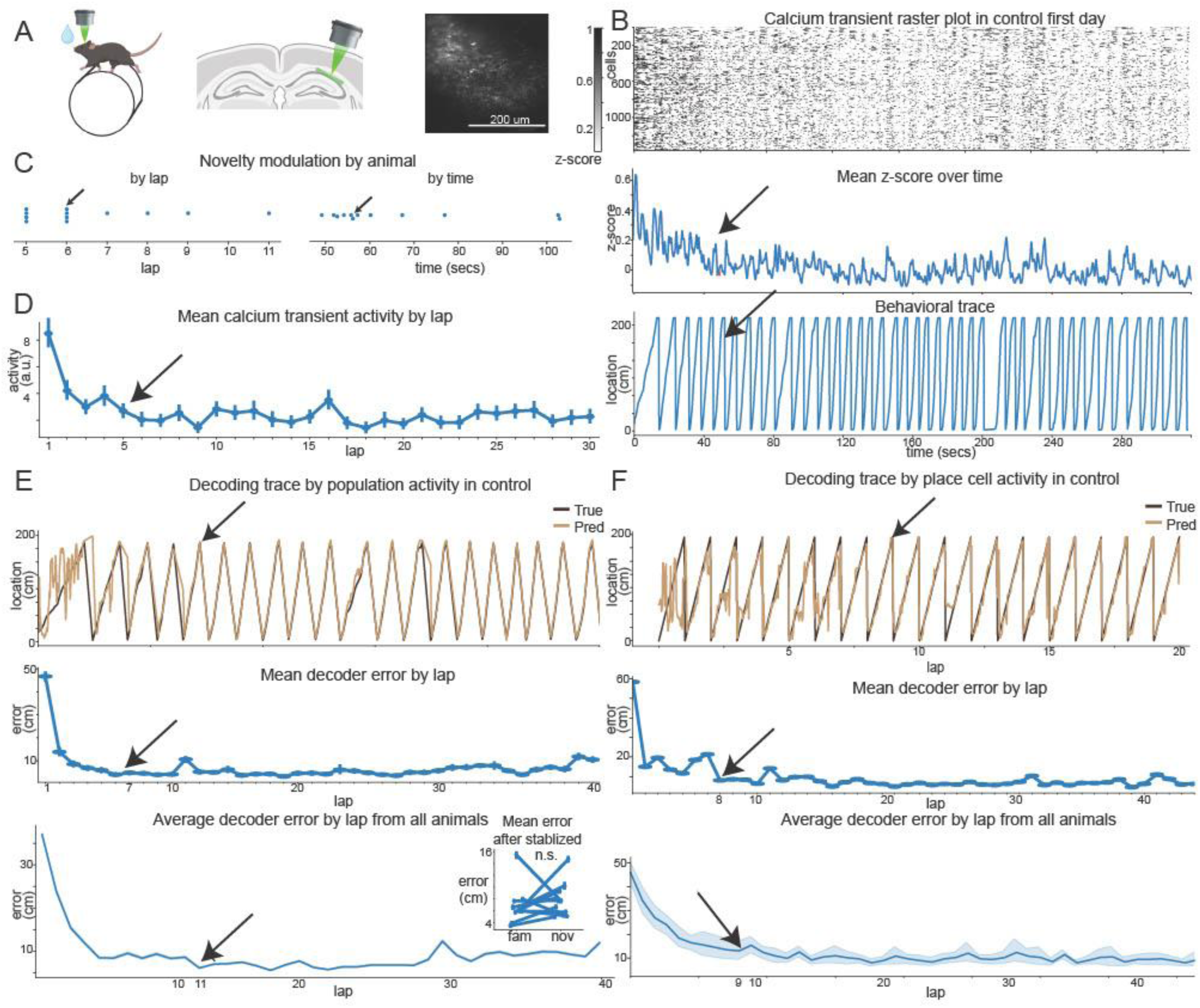
CA1 stabilization of spatial representations occurs within the first ∼10 laps in novel environments. A. Imaging setup: in vivo recording of CA1 population activity as animals navigate in novel environments. B. Z-score transformed neural activity of an example animal upon entering a novel environment. Top: Neural activity (see Methods) on control day 1, binned in ∼160 ms intervals. Middle: Mean z-score over time. The arrow marks the end of the novelty induced elevation of activity, identified using the kneedle algorithm (see methods and Supp. Fig 8) on smoothed (800 ms window) mean z-score data. Bottom: Behavioral trace of the same animal, with the arrow indicating the transition point calculated from the middle panel. C. Novelty modulation time across animals, identified by the kneedle algorithm on smoothed mean z-score data, expressed in seconds and laps. Arrows highlight the example animal from A. D. Mean calcium transient activity per lap across all cells of the example animal in A., resulted in a comparable transition lap as A. E. Decoder performance predicting location on the track trained using population activity dimension-reduced by CEBRA and a KNN regressor (see Methods). Top: Decoder-predicted (light) vs. true (dark) location for the example animal. Middle: Mean decoder error per lap for the same animal, calculated as the mean absolute difference between true and predicted location. Bottom: Mean decoder error per lap averaged across animals. The arrow marks the transition lap where decoder error plateaued, identified using the kneedle algorithm. Inset: Mean decoder error in laps after the transition lap in novel vs. familiar conditions. No significant differences were found (Wilcoxon test, p=0.45, n animals=12). F. Same as D., but using only place cell activity and Bayesian decoder (see Methods) resulting in similar transition laps as D.

## Results

We trained mice to run in a 2-meter 1D linear track virtual environment (Fig. 1a). Mice received water reward by the end of the track and were teleported back to the beginning. In the familiar environment animals were trained in, we found that animals robustly and precisely slowed down and licked at locations on the track prior to the reward site (Fig. 1b, c, green; Fig. 1d; Supp. Fig. 1). In comparison, during the initial laps in novel environments, as the animals were learning, they ran at a constant lower velocity and licked all over the track (Fig. 1b, middle; Fig. 1d right; Supp. Fig. 1). Over a few laps, animals developed faster running and precise anticipatory licking that reached a plateau close to the behavior in the familiar environment by lap 8 (Fig. 1e; Supp. Fig. 1). The development of precise anticipatory licking and running behavior reflects spatial learning as animals undergo familiarization to the environment^52^. Once precise anticipatory licking is established, we consider animals to be familiar with the novel environment.

Using two-photon microscopy, we recorded calcium transients from pyramidal cell populations in CA1_R_ as animals were navigating in novel environments (Fig. 2a). Upon immediate novel exposure, we observed elevated neural activity (Fig. 2b-d) that lasted under 2 minutes and within the first 5-10 laps (Fig. 2b-d) as animals familiarized with the environment. We trained decoders to predict location on the track during and following this learning process. CA1 decoding accuracy using population activity (all cell; Fig. 2e) and place cell firing (place cells; Fig. 2f) were initially low but progressively and rapidly improved, plateauing within ∼10 laps and matching decoder performance in the familiar environment (Fig. 2e, bottom inset). Importantly, both decoding accuracy (Fig. 2) and anticipatory licking behavior (Fig. 1) showed quick development and plateaued within the first ∼10 laps (or 2 minutes) in the novel environment. Thus, we consider this early phase in a novel environment (the first 10 laps) to be the learning phase in which familiarization to the environment is taking place. Following this early phase, we consider animals to be familiar with the novel environment.

To examine the contribution of bilateral CA3 inputs to spatial learning at the early familiarization and late familiarized phases in novel environments, we optogenetically inhibited left or right CA3 at these early and late phases while recording CA1 cell population activity. To achieve this, we injected cre-dependent opsin eOPN3 ^30^ in either left or right dorsal CA3 of Grik4-cre mice (Fig. 3a). eOPN3 selectively suppresses synaptic transmission through the Gi/o signaling pathway. It does not affect membrane voltage and therefore does not inhibit action potential firing^55^ We selected eOPN3 for this specific reason, as it allowed us to selectively inhibit either CA3L or CA3R terminals independently in CA1_R_. Grik4 mice have restricted cre expression in CA3 pyramidal cells, ^19^ although we did find minimal expression in CA2 (∼7% of CA2 cells) and dentate gyrus (∼18% of granule cells), those cells had much lower expression levels than CA3 pyramidal cells (Supp. Fig. 2). In the same animals we also expressed GCaMP6f in dorsal CA1_R_ pyramidal cells. Using two-photon microscopy, we recorded calcium transients from pyramidal cell populations in CA1_R_ and delivered light pulses to dorsal CA1_R_ through the same objective to optogenetically inhibit either left or right CA3 inputs to dorsal CA1_R_.

**Fig 3.**
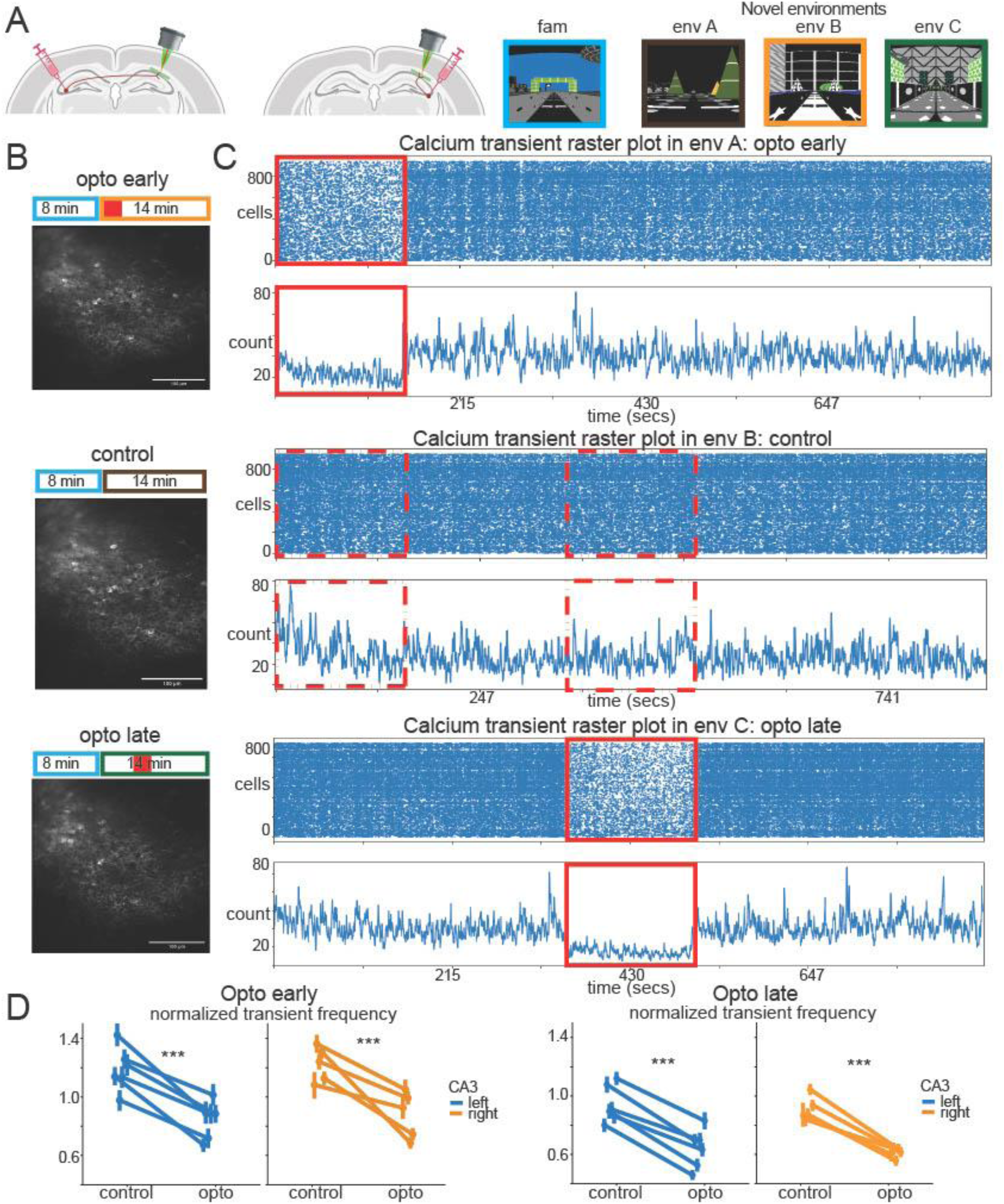
Optogenetic inhibition of left and right CA3 axons reduces CA1 pyramidal cell activity. A. Left: Viral injection protocol. GCaMP6f is expressed in CA1_R_ for imaging of CA1 pyramidal cells (green) and eOPN3 is expressed in left or right CA3 for axon inhibition in CA1_R_ (red). Right: The VR environments (familiar and 3 novel environments). B. The 3-day recording protocol. The same FOV was recorded each day (example from one mouse shown). The environments and time in the environments are indicated above the FOVs by the color-coded boxes that match the environments in A. Example behavior traces are shown below the FOVs and the opto-on periods are indicated in red. C. Example animal calcium transient raster plot from the same population of CA1 pyramidal cells in the 3 different novel environments. Each dot in the raster plot indicates a single calcium transient. Below each raster plot is the number of cells with a calcium transient per 160ms time-bin. Solid red box: opto-on periods. Dashed red box: No-opto time-matched control period. D. Opto-inhibition reduces normalized transient frequency in both opto early and late conditions in every animal. Normalized transient frequency is calculated by dividing transient frequency during opto on period (solid or dashed red box) by transient frequency during the entire time. Wilcoxon signed-rank test was performed on the same cells from the same animals in opto vs. control on each animal after correcting for multiple comparison (for 11 animals in total). p value was smaller than 0.001 for each animal in both opto early and late conditions. Same number of cells in each animal between early and late conditions=946, 666, 409, 764, 771, 761, 804, 485, 829, 1387, 926, 1085

We designed an inhibition and imaging protocol that contained two optogenetic conditions and a control condition. Each mouse experienced all 3 conditions, each one on a different day and each condition associated with a different novel environment. Also, in each mouse the same fields of view (FOVs) were tracked across all 3 conditions (Fig. 3b, c). The conditions: “Opto early” provided inhibition immediately upon entering into a novel environment and remained on for 3 minutes, “Opto late” provided inhibition after 3 minutes in a different novel environment on a different day and remained on for the subsequent 3 minutes. “Control” provided no optogenetic inhibition but the early and late phases were time-matched to the opto conditions for comparison. We a priori selected a 3-minute optogenetic inhibition for the “opto early” condition to span the period when familiarization to the novel environment is taking place (Fig. 1, 2; Supp. Fig. 1). Importantly, our opto early condition provided inhibition throughout this early learning phase. In contrast, the opto late period was applied only after familiarization had occurred, again determined a priori, i.e. once animals had developed stable and precise anticipatory licking behavior (Fig. 1d) and CA1 decoding accuracy and neural activity had stabilized (Fig. 2a-f).

Animals entered the inhibition and imaging protocol after passing behavioral criteria of running >2 laps per minute for 3 consecutive days and confirmed efficacy of opto inhibition by checking opsin expression in CA1 and testing inhibition of CA1 activity while animals are in a dark environment. On each day animals ran through the familiar environment first and then one of three novel environments paired with either the opto-early, opto-late, or the control condition (Fig. 3b and c). The combination and order of novel environments and opto conditions were counterbalanced between animals. Animals momentarily slowed down during the first few laps in a novel environment compared to laps in the familiar environment (Supp Fig. 1C), but did not show differences in running during optogenetic inhibition compared to control (Supp Fig. 1D), suggesting that any changes in neural dynamics we found were not caused by changes in running speed in response to optogenetic inhibition.

We first examined the effects of optogenetic inhibition of left and right CA3 inputs on CA1 population activity. For each animal, we binarized calcium transients from all pyramidal cells per 160 ms time-bins and counted the number of transients in each time bin (Fig. 3c). We calculated the normalized transient frequency for each cell (calculated by transient frequency during opto normalized by transient frequency across the entire session) in the opto condition and compared to the time-matched control condition in the same animal. Transient activity during stationary periods were excluded to match for behavior. All animals showed reduced transient frequency upon optogenetic inhibition (Fig. 3d), confirming that optogenetic inhibition worked on all animals.

## Right CA3 Inputs Underlie Early CA1 Spatial Refinement, Whereas Left CA3 Inputs Sustain Late Stability

Besides facilitating the firing activity of individual CA1 pyramidal cells, bilateral CA3 inputs have also shown to contribute to CA1 population spatial coding ^24,32,33^. We investigated if left and right CA3 inputs made distinct contributions to CA1_R_ spatial coding during the early and late phases following novel environment exposure. Multiple studies have identified neural manifolds within CA1 population activity that encode both task-relevant and irrelevant information using various dimensionality reduction techniques ^34–37^. To uncover low-dimensional trajectories of CA1 population activity and the effect of optogenetic inhibition of CA3 inputs, we applied CEBRA ^38^, a nonlinear dimensionality reduction algorithm that links neural dynamics to behavioral data. The resulting manifolds preserved the topology of the 1D VR environment and exhibited clear linear separation in latent space (Fig. 4a, left & middle), consistent with prior findings on structured spatial representations in CA1 ^34,35^. During optogenetic inhibition (red-coded; Fig. 4a, right), the projected manifold spread out farther from the centroid compared to the time-matched control condition (Supp. Fig. 3a, see methods), suggesting a potential disruption in the low-dimensional neural representation of the environment.

**Fig 4.**
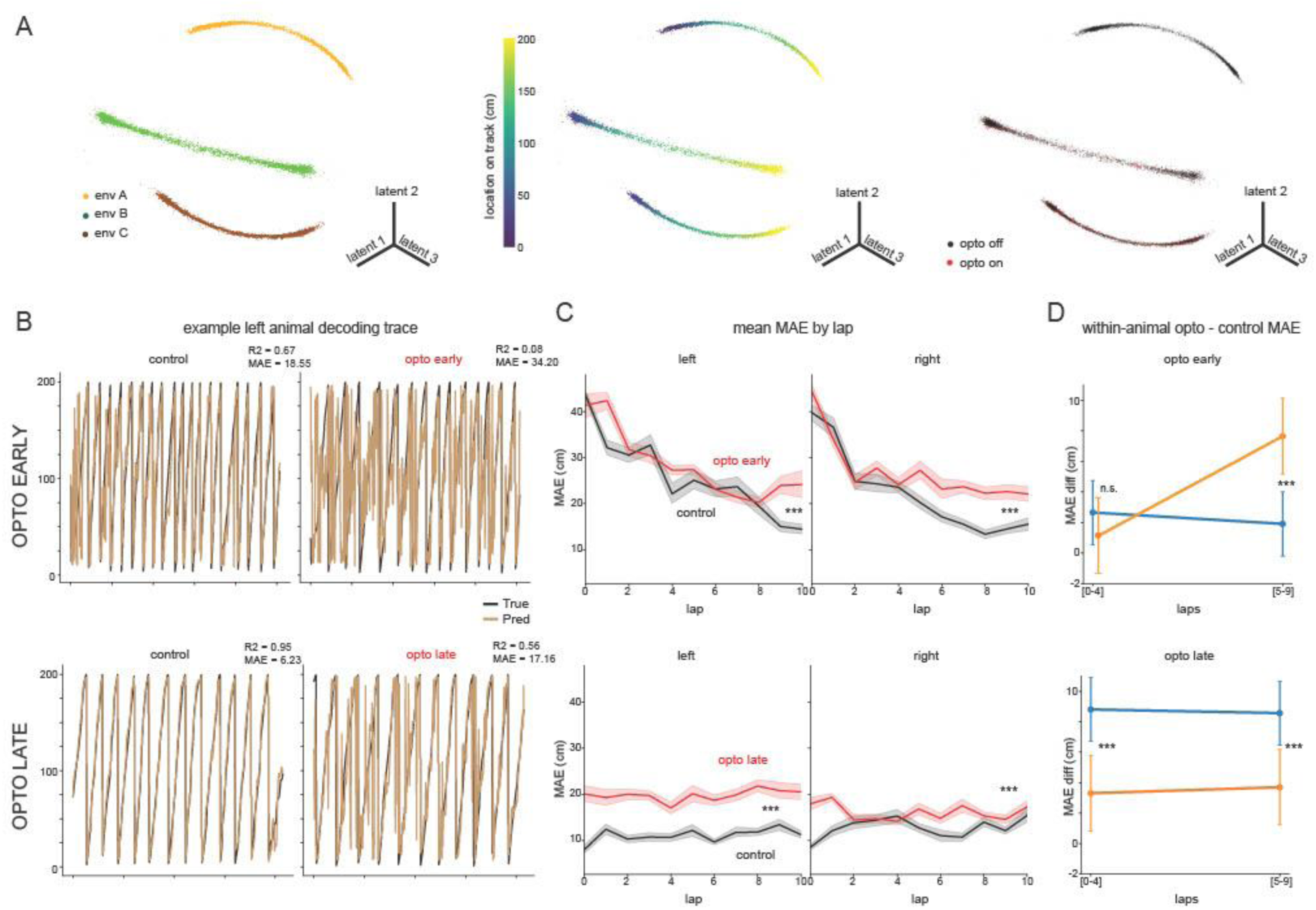
CA1 low-dimensional neural manifolds and spatial decoding reveal a dynamic contribution of left and right CA3 inputs that switches between initial learning and post-learning phases. A. Neural manifold color coded by environment (left), location on the track (middle), opto condition (right). Each point corresponds to the low dimensional representation of binarized population activity per 160ms. B. True running trajectory (dark) vs. decoded running trajectory (light) from latent embeddings in time matched control condition and opto condition during different phases. C. Decoder error per lap represented by mean absolute error (MAE) in cm averaged across animals. Both left and right opto-inhibition significantly increased MAE during opto-early (top) and opto-late (bottom) phases. Wilcoxon signed rank test comparing within animal MAEbetween opto and control: p<0.001 for all conditions after Bonferroni correction (*4) N shuffles = 500; Left n observations = 1146, right =1000 D. D. Time-matched within-animal decoder MAE difference between opto and control condition. Each data point represents MAE difference between opto and control shuffled from the same animal and the same frame grouped by lap bins. Below 0 reflects that the decoder performed worse under control. Statistical inference used a linear mixed-effects model with mouse as a random effect and fixed effects of CA3 side, opto condition, and lap bin (0–4 vs 5–9). The model showed a significant CA3 × condition × lap bin interaction (β = –6.60, p < 0.001). Predicted MAEs (95% CI): opto-early (laps 0-4) left: 2.65 [0.56, 4.74], right: 1.15 [-1.33, 3.62]; opto-early (laps 5–9) left = 1.9 [–0.2, 4.0], right = 7.6 [5.2, 10.1]. left n=6 right n=5 animals; opto-late (laps 0-4) left = 8.81 [6.72, 10.90], right = 3.3 [0.83, 5.78] (laps 5-9) left = 8.6 [6.5, 10.7], right = 3.7 [1.2, 6.2] left n=7, right n=5 animals. Total n observations=92000.

To test this, we assessed whether location on the track during opto-on periods could be accurately decoded using a latent embedding learned during opto-off periods. Under time-matched control conditions, location decoding accuracy was high (Fig. 4b, c). In the control condition, decoder performance improved rapidly over the first few traversals (Fig. 2e, f, 4c), resembling a spatial learning process in which CA1 representations emerge and refine. In contrast, in the late-phase in the control condition, decoder performance was stable across laps, suggesting CA1 representations had already stabilized with familiarity. However, optogenetic inhibition significantly impaired decoding accuracy during both early and late phases (Fig. 4c). Analysis of CA1 activity in the same task following chemogenetic inhibition of granule cells in the right hippocampus^57^ did not reveal a similar decrease in decoding accuracy, suggesting the results were not affected by possible effects on off-target granule cell labeling (Supp Fig 3)

To further examine how much impairment left and right CA3 input inhibition had on CA1 spatial representations and whether impairments were dependent on progression through the early or late phase, we compared the lap-matched within-animal decoder performance between opto and control and compared the first half to the second half of laps within each phase. We used a linear mixed-effects model with mouse as a random factor and CA3 side (left or right), condition, and lap bin as fixed factors to quantify the extent of impairment due to optogenetic inhibition between the left and the right. Interestingly, right CA3 inhibition had a significantly stronger effect than left CA3 inhibition on reducing decoder performance during the latter half of the early phase, whereas left CA3 inhibition had a significantly greater impact than right CA3 inhibition throughout the late phase (Fig. 4d). These results indicate that right CA3 inputs contribute more to the development of CA1 spatial representations by progressively driving greater spatial accuracy during early novel exposure, while left CA3 inputs maintain spatial representations once they have stabilized.

## Right CA3 inputs control early CA1 place cell dynamics whereas left CA3 inputs dominate later

To investigate how individual place cells contribute to the disruptions in population coding described above, we analyzed lap-by-lap dynamics of CA1 place cells in response to optogenetic inhibition of left and right CA3 inputs. Since inhibiting either input reduced CA1 pyramidal cell firing frequency (Fig. 3d), we assessed whether in-field firing within place fields, critical for spatial coding ^2,39^, was affected.

We first detected place cells with significant place fields based on their activity across all laps during the session. We then assessed place field reliability, a measure of how often a place cell fires in its place field across laps. Specifically, we quantified the number of laps with an in-field calcium transient within a 6-lap sliding window for each place field (Fig. 5a). During the early phase, inhibiting either left or right CA3 inputs significantly reduced place field reliability (Fig. 5b, c) but right CA3 inhibition had a greater impact than the left (Fig. 5b, inset). Similar to the effect on spatial decoding accuracy, right CA3 input inhibition prevented the development of reliable place fields. However, left CA3 inhibition more strongly affected reliability than the right during the later phase (Fig. 5c, inset).

**Fig 5.**
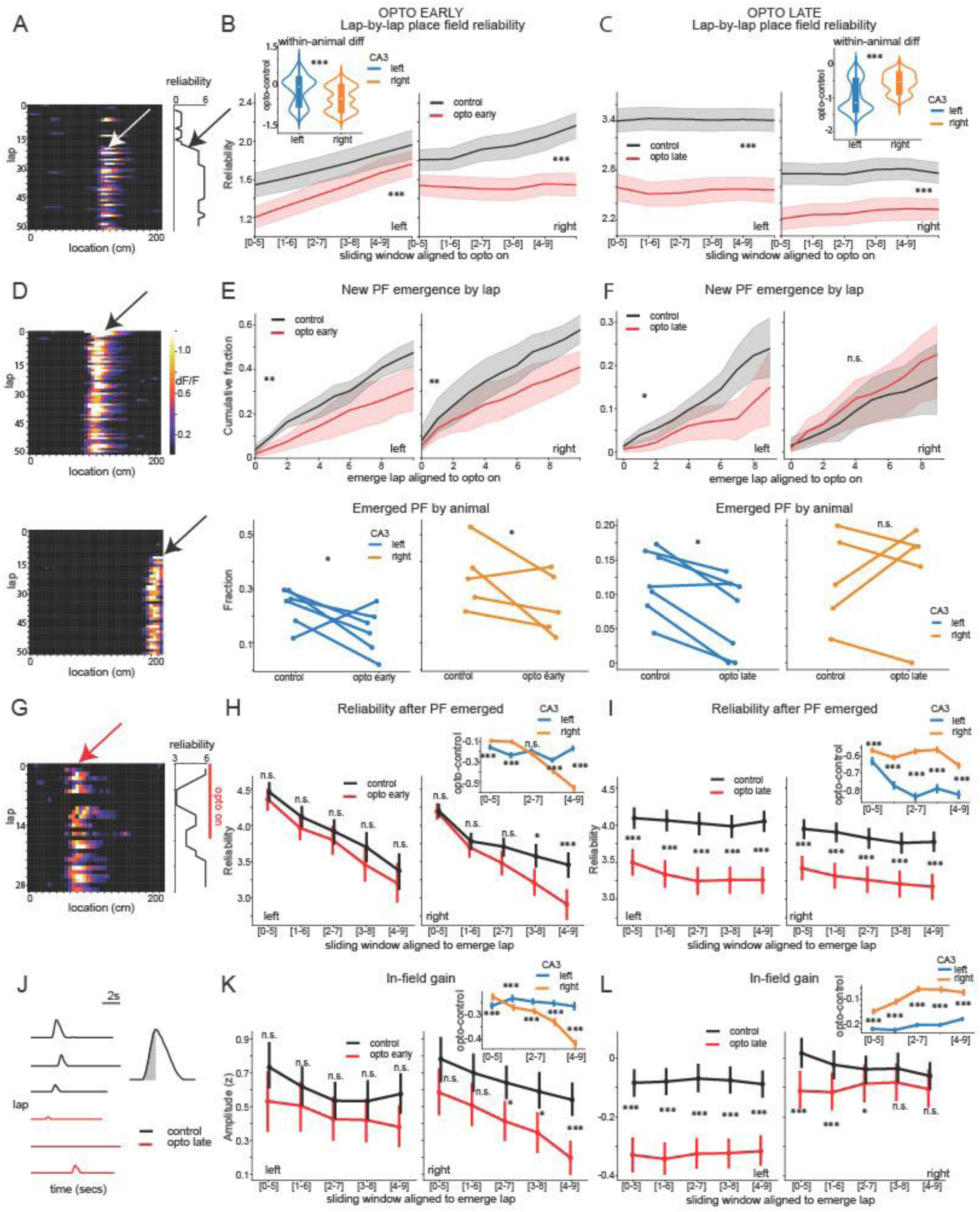
The influence of left and right CA3 inputs on the emergence, reliability, and gain of CA1 place fields during initial learning and post-learning phases. A. Example place field activity and its reliability measured over a 6 lap sliding window. 0 is inactive and 6 is always firing in-field for 6 laps. The cell only began consistently firing within-field after lap18 (pointed by the white arrow), represented by a step-like increase in reliability (black arrow). B. Average in-field firing reliability across all place fields for first 10 laps under optogenetic inhibition compared to within-animal lap-matched control. Both left and right CA3 input inhibition significantly reduced reliability (Mann-Whitney U test p<0.001, n place fields: left = 794, right=870). Inset: Bootstrapped data for within animal difference distribution. Above 0: place fields are more reliable in opto versus control. Below 0: more reliable in control. Right CA3 inputs inhibition reduced place field reliability significantly more than left CA3 input inhibition during opto-early (Mann-Whitney U test p<0.001, n shuffles=500, n observations left=18000, right=15000). C. Same as B for opto-late. Both left and right CA3 inputs inhibition significantly reduced in-field firing reliability (Mann-Whitney U test p<0.001 between opto and control for both left and right, n place fields: left=639, right=704). Inset: Left CA3 inputs inhibition significantly reduced place fields reliability more than right CA3 input inhibition in opto-late (Mann-Whitney U test p<0.001, n shuffles=500, n observations left=17500, right=12500). D. Two example place fields; one emerged on lap 1 (top) and one on lap 18 (bottom). Emergence lap is defined as the first lap in which 3 out of 6 subsequent laps fired. E. Top: Cumulative histogram showing the proportion of place fields that have emerged as a function of laps during opto-early (red) versus control (black). Both left and right CA3 inhibition significantly delayed the emergence of CA1 place fields. Wilcoxon Signed-Rank Test left: p=0.026, right: p=0.003; n laps left=60, right=50. Bottom: Within-animal comparison of fraction of place fields emerged over laps under opto-inhibition and control. Both left and right CA3 inhibition significantly decreased the number of fields that emerged (linear mixed effects model, main effect of opto p=0.005, interaction term opto x CA3 p=0.97, CA3 p=0.062, n animals=11). F. Same as E for comparing opto-later (red) with control (black). Top: Only left CA3 inhibition significantly delayed the emergence of place fields. Wilcoxon Signed-Rank Test left: p=0.014; right: p=0.143; n laps left=70, right=50. Bottom: Only left CA3 inhibition significantly reduced the fraction of fields formed under opto compared to lap-matched control. (linear mixed effects model, main effect of opto p=0.004 and interaction term opto x CA3 p=0.017, CA3 p=0.923. Wilcoxon signed-rank test left: p=0.031; right: p=0.19, n animals=12). G. Example place field showing emergence lap (red arrow) and reliability (right). H. Reliability of place fields after emergence for opto early (red) and control (black). Left CA3 input inhibition did not affect place fields reliability: 5 lap windows: p=1.2, 0.8, 1.5, 0.35, 1.7; n place fields opto vs. control = 308 & 398, 268 & 360, 225 & 314, 186 & 261, 132 & 174, respectively. Right CA3 input inhibition significantly reduced reliability in the last two sliding windows: 5 lap windows: p=4.6, 2.1, 0.11, 0.011, <0.001, n place fields in opto vs. control = 432 & 725, 393 & 671, 347 & 612, 304 & 549, 236 & 435, respectively. Inset: Bootstrapped distribution within-animal difference in mean reliability between opto and control (see methods). Right CA3 inhibition reduced reliability significantly more than left in the last two sliding windows. 5 lap windows from left to right: p<0.001, <0.001, 0.15, <0.001, <0.001, n shuffles=500, n observations left=3000, right=2500 for each sliding window. All tests used Mann-Whitney U test with Bonferroni correction (*5). I. Same as H for opto late. Both left and right CA3 inhibition significantly reduced place field reliability: p<0.001 for all sliding windows; Left n place fields in opto vs. control = 515 & 668, 514 & 666, 509 & 661, 506 & 654, 504 & 650; Right n place fields in opto vs. control = 466 & 643, 463 & 636, 454 & 625, 447 & 614, 442 & 605. Inset: Same as H in opto late. Left CA3 inhibition reduced reliability significantly more than right CA3 inhibition (p<0.001 for all sliding windows after Bonferroni correction *5, n shuffles = 500, n observations left = 3500, right = 2500). J. Calcium transients from an example place field without optogenetic inhibition (black) and with optogenetic inhibition (red). Transient magnitude is calculated as the area under the curve from rise to peak (shaded area). K. Left: Normalized in-field transient magnitudes in control vs. opto early from all transients pooled together across animals within each 6-lap sliding window. Within each sliding window, all tests used Mann-Whitney U test with Bonferroni correction (*5) on in-field transients, left CA3 input inhibition did not significantly reduce in-field transient magnitudes p=0.1, 0.25, 0.56, 0.5, 0.1 respectively. N transients for opto vs. control in each sliding window = 105 & 140, 130 & 152, 143 & 174, 156 & 191, 166 & 210, respectively. Right CA3 input inhibition gradually became significant over laps, p=0.75, 0.35, 0.03, 0.011, <0.001. N transients for opto vs. control in each sliding window = 136 & 245, 147 & 265, 163 & 292, 177 & 310, 188 & 330, respectively. Inset: Bootstrapped distribution within-animal difference in mean transient amplitude between opto and control throughout each sliding window (see methods). Except for the first sliding window, where left CA3 inputs inhibition significantly decreased in-field gain more than right CA3 inputs (two sample ttest n shuffles = 500, left observations =3000, right observations = 3000, p<0.001 after Bonferroni correction *5), right CA3 inputs inhibition significantly decreased in-field gain more than left CA3 inputs for the rest of the sliding windows (same number of shuffles and observations, p<0.001 after Bonferroni correction for each sliding window). L. Right: Same as K for opto late. Both left and right CA3 inhibition had sliding windows where transient magnitudes were significantly reduced Left: Left CA3 input inhibition significantly reduced in-field gain in all sliding windows under Mann-Whitney U test with Bonferroni correction (*5) p<0.001 for all sliding windows, N transients for opto vs. Control in each sliding window = 426 & 592, 422 & 603, 433 & 609, 445 & 615, 449 & 621 respectively. Right: Right CA3 inputs inhibition reduced in-field transient magnitudes significantly in the first three sliding windows, but were insignificant in the rest of the two sliding windows. Mann-whitney U test with Bonferroni correction (*5) p <0.001, <0.001, 0.035, 0.065, 0.1. N transient for opto vs. control in sliding windows = 412 & 554, 420 & 564, 421 & 578, 429 & 583, 432 & 585. Inset: Same as K for opto late. Left CA3 inhibition significantly reduced in-field gain more than right CA3 inputs for all sliding windows (two sample t-test, p<0.001 for all 5 sliding windows after Bonferroni correction *5, n shuffles=500, left n observations = 3500, right observations=3000).

Reliability could be influenced by both newly emerging place fields—characterized by abrupt, step-like increases in reliability as they transition from absent or sporadic firing to consistent in-field activity (Fig. 5d)—and by the lap-lap reliability of already established place fields (Fig. 5g). To disentangle these contributions, we directly tested whether left versus right CA3 inhibition differentially affected CA1 place field emergence.

We defined the emergence lap as the first lap in which subsequent in-field firing became reliable, requiring at least 3 out of 6 consecutive laps to contain in-field activity. Upon novel environment exposure, CA1 place fields typically emerge rapidly within the first few laps ^7^. Consistent with this, many CA1 place fields formed quickly under control conditions (Fig. 5e, black lines). However, optogenetic inhibition of either left or right CA3 inputs during initial novel exposure significantly delayed place field emergence (Fig. 5e, upper panel) and reduced the total number of fields formed (Fig. 5e, lower panel), with no significant difference between left and right CA3 inhibition (Supp Fig. 5b) and this effect was not due to our definition of emergence (Supp Fig. 5a, Supp Fig. 5b). This indicates that the stronger impact of right CA3 inhibition on place field reliability is not due to a greater disruption of place field emergence.

To further investigate the impact of CA3 inhibition on place field reliability, we examined whether already emerged place fields became less reliable over successive laps under inhibition (Fig. 5g). Following the formation of place fields, reliability gradually declined in the control group (Fig. 5h, black lines), potentially due to reported inhibitory dynamics in CA1 that show initially low inhibition in novel environments that gradually increase during the early phase ^12,13^. We found that during the early phase, inhibiting the right CA3 inputs reduced place field reliability, with this effect becoming more pronounced over laps and reaching statistical significance towards the end of the early phase (Fig. 5h). In contrast, left CA3 inhibition had no significant effect on place field reliability at any point during the early phase (Fig. 5h). These results indicate that the stronger impact of right CA3 inhibition on place field reliability is specifically due to decreased reliability of emerged place fields rather than a delay in their emergence.

In contrast, during the later phase, both left and right CA3 inhibition significantly reduced place field reliability (Fig. 5i), with left CA3 inhibition having a significantly stronger effect than right CA3 inhibition throughout the phase (Fig. 5i, inset). Additionally, left CA3 inhibition during the later phase also impaired new place field emergence, whereas right CA3 inhibition did not (Fig. 5f). These findings indicate that while both left and right CA3 inputs are necessary for the rapid emergence of CA1 place fields during initial novel exposure, right CA3 primarily regulates place field reliability early on, whereas left CA3 plays a greater role in maintaining reliability later and continues to drive new place field emergence during the later phase.

The reliability analysis assessed whether a place cell fired in its place field on a given lap (a binary measure) but did not account for variations in firing rate within place fields. Place cells modulate their firing rates in response to contextual changes (rate remapping), reflecting both task-relevant and irrelevant cognitive processes ^40,41^. As a result, CA1 place field gain carries important information that can influence downstream circuits ^42,43^. To examine whether left and right CA3 inputs differentially regulate place field gain across memory phases, we quantified the area under the curve (AUC) during the rise-to-peak period of calcium transients on each lap as a proxy for firing rate (Fig. 5j). To facilitate comparisons across cells and experimental conditions, we z-score transformed within-field transient amplitudes, focusing on relative changes rather than absolute values.

Optogenetic inhibition of left and right CA3 inputs on in-field transient amplitudes were dependent on the phase of learning. During opto early, right CA3 input inhibition had increasingly larger effects in reducing transient amplitudes over learning, especially compared to the left CA3 inputs (Fig 5k). This again suggests that right CA3 inputs are essential for the development of CA1 spatial properties during the early phase. In addition, when we measured the effect of CA3 input inhibition as a function of calcium transient amplitude, we found right CA3 inputs (and left) had a bigger impact on reducing the activity of higher amplitude transients (Supp Fig 6a). This suggests right CA3 inputs have their biggest influence on high frequency firing (possibly burst firing) within place fields during initial experience in a novel environment.

During opto late, the effects of optogenetic inhibition on in-field transient amplitudes were reversed, with left CA3 input inhibition having a consistently larger effect in reducing in-field transient amplitudes throughout all sliding windows (Fig 5l). Unlike during the early phase, this effect was not through a selective inhibition on high amplitude transients. Instead, left CA3 input inhibition reduced the amplitude of all transients (Supp Fig. 6a). Interestingly, right CA3 input inhibition had a bigger impact on the lower amplitude in-field transients, directly opposite to its effects during opto early. These results suggest that right and left CA3 inputs control CA1 place field gain and influence distinct modes of firing to different extents dependent on the phase of learning, with right inputs dominating early and left inputs dominating later.

## Lateralized CA3 axon dynamics underlie phase-specific contributions to CA1 during initial learning and post-learning phases

To investigate whether differences in left and right CA3 input dynamics underlie their distinct roles in CA1 during early and late phases of novel environment exposure, we recorded the activity of left or right CA3 axons in right CA1. Using Grik4-Cre mice, we injected Cre-dependent axon-GCaMP6s into either left or right CA3 and performed in vivo two-photon calcium imaging of CA3 axons in the stratum oriens of CA1 during spatial navigation in a familiar environment and two novel environments over two days (Fig. 6a). All subsequent analyses focused on the first exposure to the two novel environments combined.

**Fig 6.**
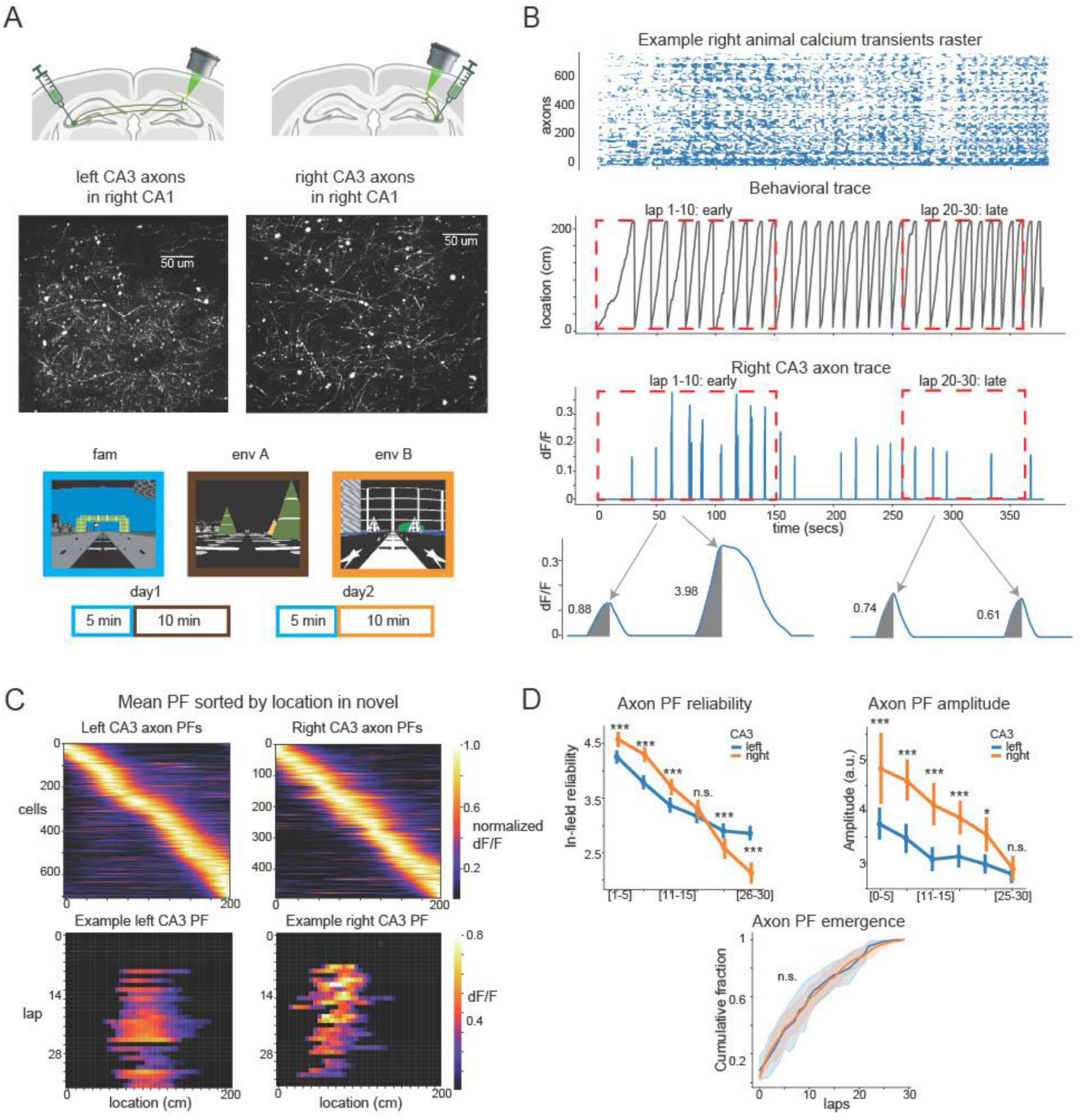
Recordings of CA3 axons reveal a switch from stronger right CA3 axon activity during the early phase to stronger left CA3 axon activity during the later phase. A. Left: Cre-dependent axon gcamp6s was injected to Grik4-cre mice in left/right CA3. Middle: Example FOV of axon population recording of left and right CA3 axons in right CA1 at S.O. Right: Example axon calcium activity over time after preprocessing. B. Top: Example calcium transient raster of all axons in env A. Middle: Behavioral trace. First 10 laps and lap 20-30 are highlighted in red box. Bottom: Example right CA3 axon calcium transient over time. C. Top: Place fields detected from axon recordings pooled together by CA3 sorted by location on the track. Bottom: Example individual place fields from left and right CA3 axons. D. In-field axon transient reliability, in-field gain, and field emergence measured from using the same methods as Fig. 5 binned every 5 laps over experience in novel environments. Top left: Right CA3 axon place fields had significantly higher reliability than left CA3 axon place fields in the first 3 lap bins (before lap 15). No significant differences between left and right in lap bin from 15 to 20 laps. In contrast, in lap bins after lap 20, left CA3 axons place fields had significantly higher reliability than right CA3 place fields. (For each lap bin, p<0.001, <0.001, <0.001, 0.2, <0.001, <0.001 Mann Whitney U test after Bonferroni correction. N place fields for each lap bin, left= 448, 721, 964, 964, 964; right= 286, 455, 664, 664, 664). Top right: Right CA3 axon place fields exhibited higher transient amplitudes than left CA3 during the first four lap bins, but right CA3 amplitudes declined faster over time, with left and right CA3 became insignificantly different in the last lap bin. (For each lap bin, p<0.001, <0.001, <0.001, <0.001, 0.02, 3.0 Mann Whitney U test after Bonferroni correction. N place fields for each lap bin, left= 425, 635, 747, 831, 879, 893; right= 187, 375, 468, 528, 531, 476). Bottom: Emergence: No significant differences were found between left and right CA3 axon place fields in emergence dynamics. Rank sum tests were done on the cumulative percentage of place fields that have emerged by the lap for the first 30 laps pooled from left and right animals. N=120 laps from 12 animals for both left and right. P=0.7

We first analyzed the spatial coding properties of individual CA3 axons from the left and right and found axon place fields that tiled the entire track (Fig. 6c). The emergence dynamics of these place fields were the same in the left and right axons (Fig. 6d) and their place field properties were similar (although some small differences were observed; Supp Fig. 7c). To examine how left and right CA3 inputs may contribute to CA1 place field features (Fig. 6d), we quantified axon place field reliability and gain using the same methods used for CA1 place cells. Specifically, axon transients were detected following the approach used in the *subprep* package ^44^, and transient magnitude was quantified using the area under the curve (AUC) during the rise-to-peak period as a proxy for spiking activity (Fig. 6b). We found that during the initial novel exposure, right axon place fields had significantly larger in-field gain and higher reliability, together providing a stronger, higher fidelity spatial input than left CA3 inputs (Fig. 6d). However, following familiarization to the environment, while both left and right CA3 spatial inputs diminished in reliability and gain, right CA3 inputs had a larger decline, which led to a shift to left CA3 inputs as the more reliable spatial input to CA1 (Fig. 6d). This shift, at least in part, likely underlies the transition from right CA3 dominance in controlling the refinement of CA1 spatial representations during early experience in a novel environment to left CA3 dominance following familiarization when representations are relatively stable.

## Discussion

Our results reveal that left and right CA3 inputs contribute to right CA1 spatial representations in distinct, time-dependent ways during familiarization to a novel environment – a form of incidental learning. Initially, when animals are exposed to a novel environment, both inputs are required for the rapid emergence of CA1 place fields. However, once these fields form, right CA3 inputs predominantly enhance early spatial coding by boosting the progression of place field reliability, amplitude, and overall decoding accuracy. Once the environment becomes familiar, left CA3 inputs assume a more dominant role, maintaining stable spatial representations by preserving field reliability and gain and by supporting the emergence of new fields. This functional transition is mirrored by axonal recordings in CA1: right CA3 axons show greater place-field activity and reliability early, whereas left CA3 axons become more reliable following familairization. Together, these findings demonstrate a dynamic, temporally structured division of labor between left and right CA3 inputs in shaping CA1 spatial maps during and following familiarization to a novel environment.

Manifold analysis revealed that during initial exploration, inhibiting right CA3 inputs significantly reduced the development of spatial decoding accuracy, indicating that right CA3 inputs are crucial for rapidly establishing a high-fidelity spatial map in right CA1. Since CA1 is the primary hippocampal output, its accurate spatial representations during novel exploration and memory formation likely play a critical role in driving synaptic plasticity in downstream brain regions ^2,39,43^. At the single cell level, high-gain, reliable place fields may promote robust synaptic plasticity at CA1 output synapses, supporting the formation of memory engrams beyond the hippocampus^42,46^. Disrupting the development of these dynamics through right CA3 inhibition results in less consistent CA1 output (lower gain, unreliable place fields), leading downstream neurons to receive variable or misaligned inputs that could impair memory formation. Right CA3 inputs to right CA1 may therefore be critical in this process in the right hemisphere.

Following familiarization once animals precisely predict the reward location in the novel environment, the role of CA3 inputs shifts. During this later phase, left CA3 inputs assume greater control, as inhibition of left CA3 leads to a more pronounced reduction in spatial decoding accuracy. This suggests that left CA3 is critical for refining and maintaining CA1 spatial maps once initial memory formation has occurred and the hippocampus operates in a memory retrieval and updating mode (familiarity requires memory retrieval) ^23,25^. While CA1 representations naturally drift over time – a process thought to support memory updating – excessive drift may impair memory retrieval ^47–51^. Our findings indicate that left CA3 inputs support stability in the CA1 population code by maintaining reliable, high-gain place fields, which is essential for ensuring that downstream neurons receive a consistent hippocampal output during memory retrieval. Simultaneously, left CA3 also promotes the emergence of new place fields, allowing for ongoing memory updating (drift) as new experiences are integrated.

Two-photon calcium imaging of CA3 axons in the right CA1 revealed that right CA3 axons exhibit elevated activity within axon place fields that are more spatially reliable (fire more often in their field) during the early learning phase of novel environment exposure, precisely when CA1 place fields rapidly emerge with high gain and reliability. It is likely that these CA1 place cell dynamics are therefore partly inherited by right CA3 inputs during this early phase.

Interesting, we found that CA3 inputs during the early phase almost exclusively drive high amplitude calcium transients. These high amplitude transients reflect high frequency spiking and likely often reflect burst firing^58^. Bursts are associated with behavioral timescale synaptic plasticity (BTSP) events which powerfully and abruptly shape place field dynamics^14,15^. They have also been shown to occur with higher probability during the initial experience in novel environments^16,17^. We propose that right CA3 inputs influence the progression of spatial map refinement in part by driving burst firing that increases the probability of BTSP events during the early phase in a novel environment.

Interestingly, there are differences in the sensitivity to plasticity inducing events at CA1 synapses receiving input from left or right CA3. Synapses from Left CA3 inputs onto bilateral CA1 pyramidal cells contain more GluN2B-enriched NMDARs, which facilitate synaptic potentiation, whereas right CA3 synapses have a higher density of AMPARs, supporting stronger baseline transmission ^22,28^. Electrophysiological studies suggest that LTP is induced more readily at left CA3 to bilateral CA1 synapses with certain stimulation protocols, while right CA3 to bilateral CA1 synapses do not undergo LTP under the same conditions^21,23^. This is not to suggest that right CA3 inputs are not plastic, but it does show the origin of CA3 input determines the synaptic type and the sensitivity of those synapses to plasticity inducing events. It remains to be seen whether BTSP occurs at right or left CA3-CA1 synapses, or both, or whether one type is more sensitive than the other.

Furthermore, these findings suggest that the functional impact of CA3 inputs on CA1 is determined primarily by the hemispheric origin of the CA3 neurons rather than by whether the inputs arrive ipsilaterally or contralaterally. In other words, both left and right CA1 respond similarly to left CA3 inputs and similarly to right CA3 inputs, despite the opposite laterality of the projections. We therefore hypothesize that the findings reported in this paper would be mirrored in left CA1, even though the right CA3 inputs would be contralateral and the left inputs ipsilateral.

Early novelty creates a transient window of heightened hippocampal plasticity, during which mechanisms such as dendritic spikes^12^, BTSP^16,17^, and neuromodulatory facilitation^56,65^ are engaged with high probability. These processes may enable LTP induction even from right CA3 inputs, accelerating the refinement of spatial representations during early experience. We propose that this extreme plasticity window that leads to new spatial maps is uniquely triggered by exposure to novel environments. Other forms of hippocampal-dependent learning typically induce more subtle shifts in spatial maps^59,60^. This transient plastic period likely enhances not only the formation of spatial maps but also other hippocampal-dependent learning occurring within it^8,68^. We show here that right CA3 inputs are more influential in shaping the development of new maps withing the plastic period, likely engaging plasticity-inducing mechanisms. Once spatial representations stabilize, this plastic window likely closes, and left CA3 inputs become more influential, supporting slower, sustained learning beyond the initial phase.

This framework reconciles our findings with prior studies reporting left CA3 dominance in hippocampus-dependent learning tasks^61–64^. These studies used continuous inhibition of CA3 across multi-day training, encompassing both the early novelty-driven and later stabilized phases. Given the extended training durations and task complexity, it is likely that most learning occurred after novelty-driven plasticity had subsided, explaining why left CA3 inhibition produced greater behavioral deficits. Thus, our data suggest a dynamic hemispheric specialization: right CA3 drives rapid learning during a transient plastic period, while left CA3 dominates during prolonged, slower forms of learning.

**Supp Fig 1.**
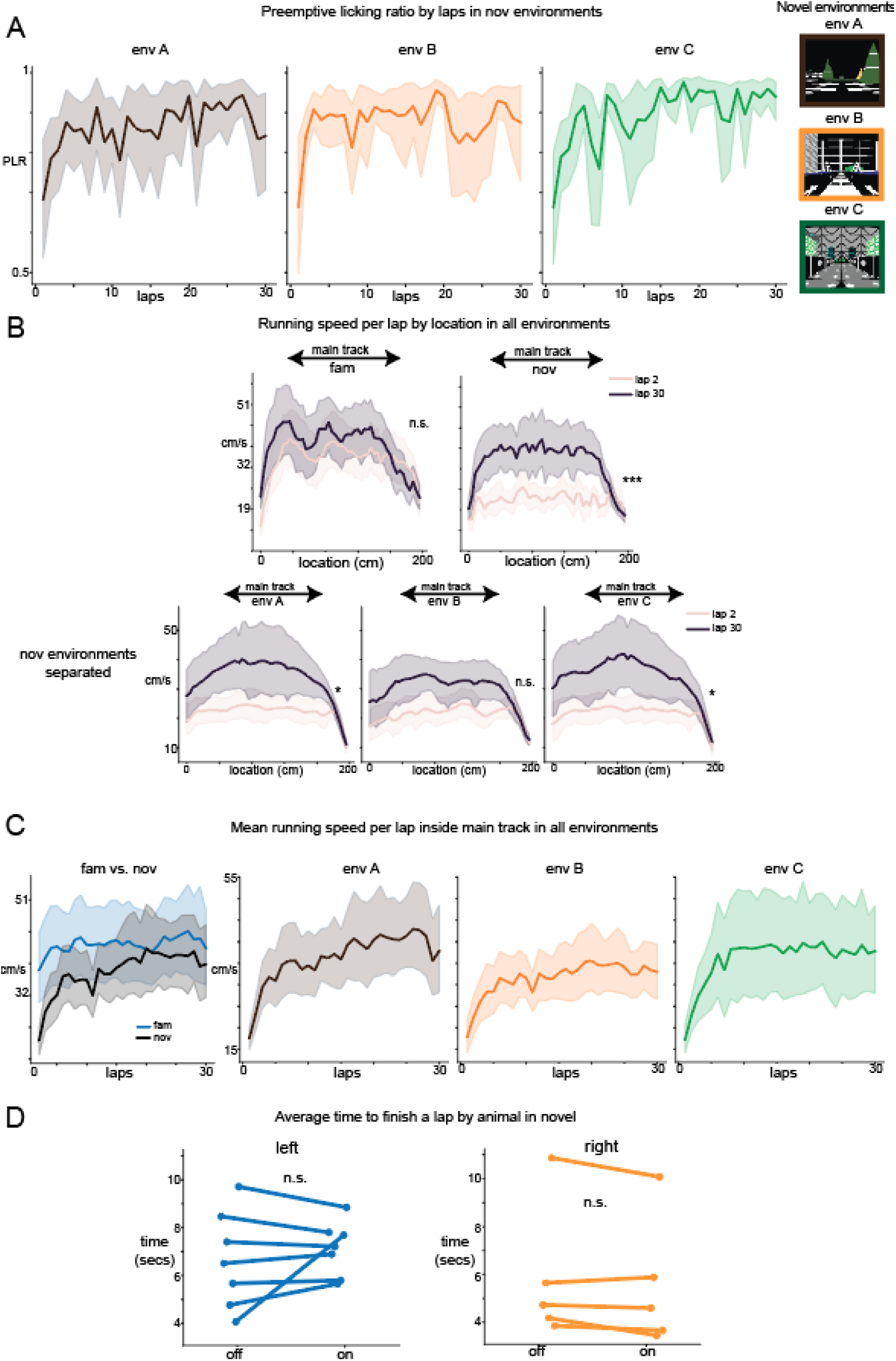
Animal behavior in familiar and novel environments. A. Preemptive licking ratio by laps separated in 3 different novel environments averaged across all animals (N=12). B. Top: Running speed from all animals (n=12) across all locations before rewards within lap 2 and lap 30 in familiar and 3 novel environments combined. Main track region here included track locations after consummatory licks and before pre-reward zone. Average velocity in lap 2 and lap 30 was not significantly different in familiar environment (Wilcoxon test p=0.23 after Bonferroni correction, n=12), but significantly different in novel environment (Wilcoxon test p<0.001 after Bonferroni correction, n=12). Bottom: Novel environments separated. Wilcoxon test p in three environments p=0.03, 0.06, 0.01 after Bonferroni correction, n=12. C. Running speed averaged across main track region over each lap in all environments. D. Average time elapsed (in seconds) for running one lap under optogenetic inhibition and before turning on opto across all mice. CA3 Optogenetic inhibition did not alter the mean running speed of animals in novel environments. Frames recorded during and after reward zone were not included. Wilcoxon test between opto off and on, left: p=0.69, n=7; Right: p=0.31, n=5. Between off and on pooled between all animals: Wilcoxon test p=0.79, n=12.

**Supp Fig 2.**
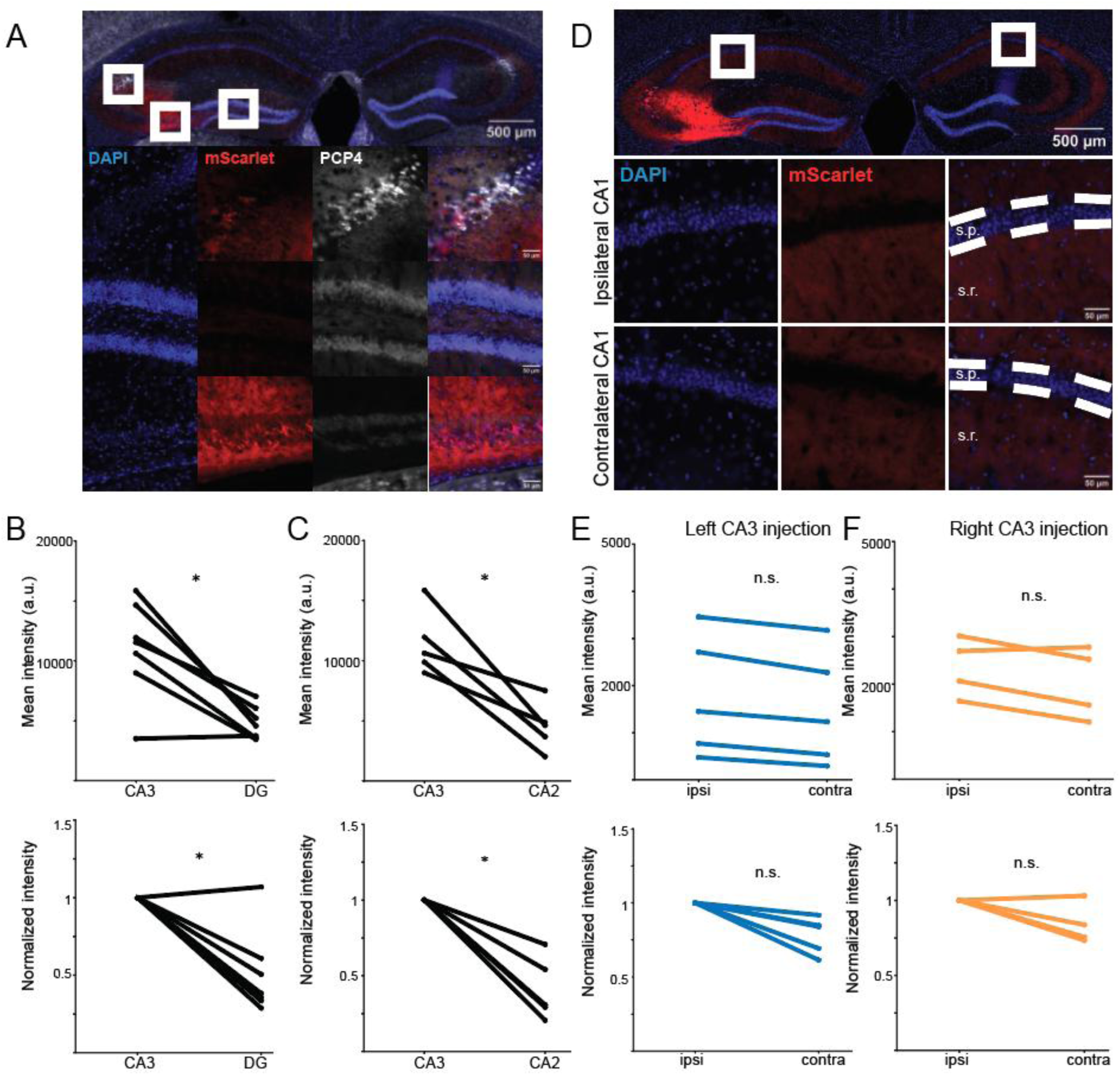
There is limited off-target eOPN3 expression in DG granule cells and CA2 pyramidal cells compared to CA3 somas. A. (Top) Representative immunofluorescent image of the hippocampus from a Grik4-Cre mouse injected with eOPN3-mScarlet into the left CA3. Magnified view of CA2 (middle row) and DG (bottom row) from the left and right white square in the top image, respectively. Shows labeling of cell nuclei with DAPI (blue), the opsin fluorescent tag mScarlet (red), and CA2 pyramidal cells with PCP4 (white). B. (Top) Mean intensity of mScarlet within CA3 pyramidal cells and DG granule cells within the injected hemisphere. Each dot represents the mean intensity calculated from three 30 μm tissue sections obtained from a single mouse (n = 7). (Bottom) Normalized intensity of mScarlet within DG granule cells. Mean values are normalized to the intensity in CA3 pyramidal cells for each mouse (n = 7). One-sided Wilcoxon signed-rank test (stat = 27.0 p = 0.016) C. (Top) Mean intensity of mScarlet within CA3 and CA2 pyramidal cells within the injected hemisphere. Each dotrepresents the mean intensity calculated as described in (B) (n = 5). (Bottom) Normalized intensity of mScarlet within CA3 and CA2 pyramidal cells. Mean values are normalized to the intensity in CA3 pyramidal cells for each mouse (n = 5). One-sided Wilcoxon signed-rank test (stat = 15.0 p = 0.031). D. (Top) Representative immunofluorescent image of the hippocampus from a Grik4-Cre mouse injected with eOPN3-mScarlet into the left CA3. Magnified view of the CA1 ipsilateral (middle row) and contralateral (bottom row) to the injection taken from the left and right white square in the top image, respectively. Cell nuclei are labeled with DAPI (blue) and the opsin with the fluorescent tag mScarlet (red). The white dashed lines mark the stratum pyramidale layer (s.p., cell bodies) and the region below the stratum radiatum (s.r., axon layer). E. (Top) Mean intensity of mScarlet in the stratum radiatum for mice injected with eOPN3-mScarlet into the left CA3. Each dot represents the mean intensity calculated from three 30 µm tissue sections obtained from a single mouse. (Bottom) Normalized intensity of mScarlet in the stratum radiatum for mice injected with eOPN3-mScarlet into the left CA3. Mean values are normalized to the intensity on the side ipsilateral to the injection (n = 5). Two-sided Wilcoxon signed-rank test (stat = 0 p = 0.0625) F. (Top) Mean intensity of mScarlet in the stratum radiatum for mice injected with eOPN3-mScarlet into the right CA3. Each dot represents the mean intensity calculated as in (B). (Bottom) Normalized intensity of mScarlet in the stratum radiatum for mice injected with eOPN3-mScarlet into the left CA3 (n = 4). Two-sided Wilcoxon signed-rank test (stat = 1.0 p = 0.25). (Top) Mean intensity of mScarlet in the stratum radiatum for all mice injected with eOPN3-mScarlet. Each dot represents the mean intensity calculated as in (B). (Bottom) Normalized intensity of mScarlet in the stratum radiatum for mice injected with eOPN3-mScarlet into the right CA3 (n = 9). Two-sided Wilcoxon signed-rank test (stat = 1.0 p = 0.0078)

**Supp Fig. 3.**
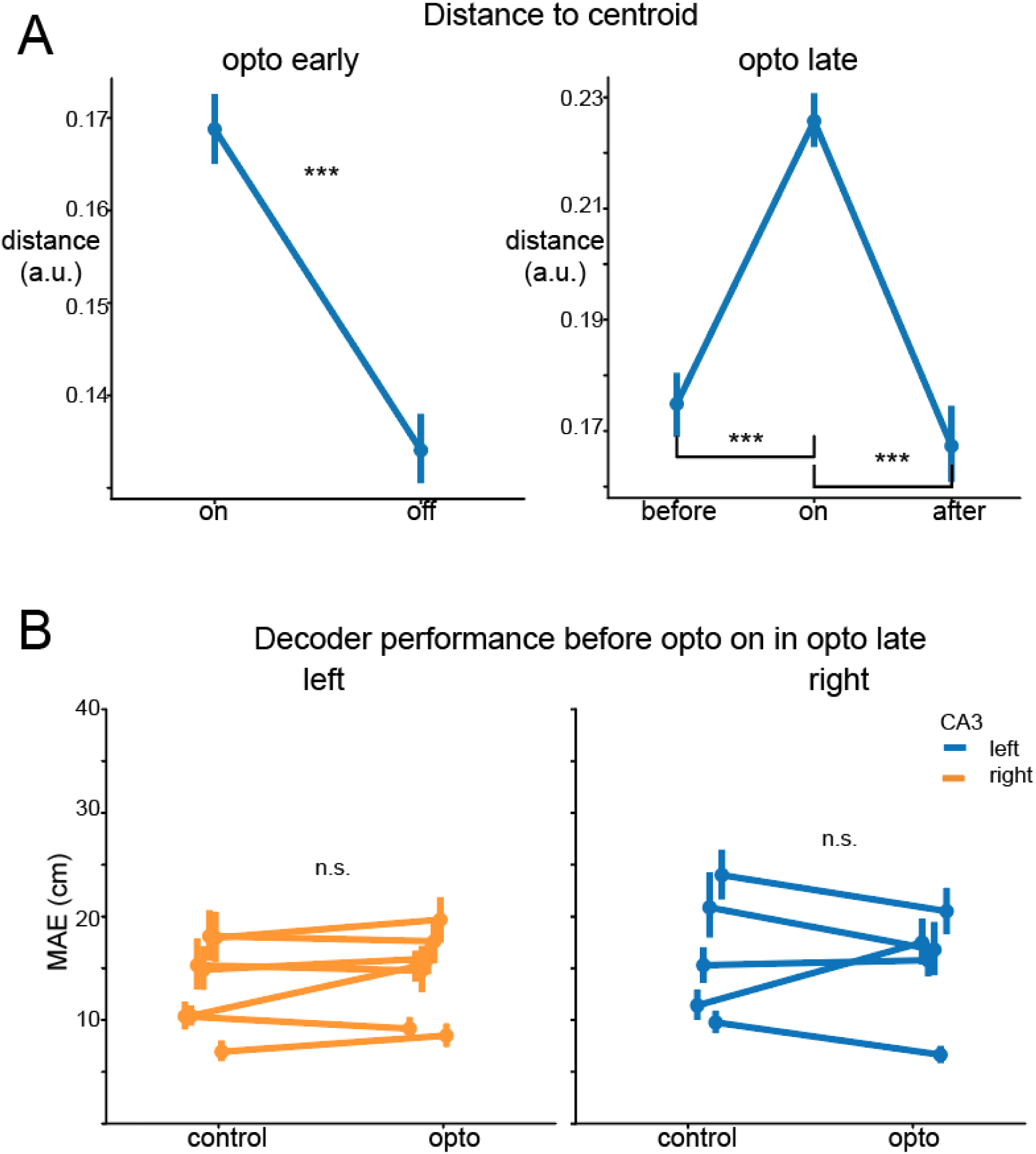
Right optogenetic inhibition significantly increased the distance to centroid in the same example animal as Fig4, representing a less stereotypical neural activity to location on the track under opto. A. Distance to centroid in opto early and opto late. Same example animal as (Fig4. a). See methods for calculating distance to centroid. Left: Optogenetic inhibition significantly increased distance to centroid than off in opto early. (Mann whitney u stats= 152935071, p<0.001). Right: Turning on optogenetic inhibition significantly increased distance to centroid than before and after. (Before vs. on: p<0.001; after vs. on: p<0.001). B. Decoder performance before turning on optogenetic inhibition in opto late and control condition. No significant difference in decoder performance between opto and control were found in both left and right CA3 (Wilcoxon signed rank test comparing within animal difference after Bonferroni correction (*2) left: p=0.16, n observations=1470; right: p=0.5, n observations=1000). Decoder performance trained using the same protocol (see methods for details) tested in 5 laps before opto on in opto late condition.

**Supp Fig 4.**
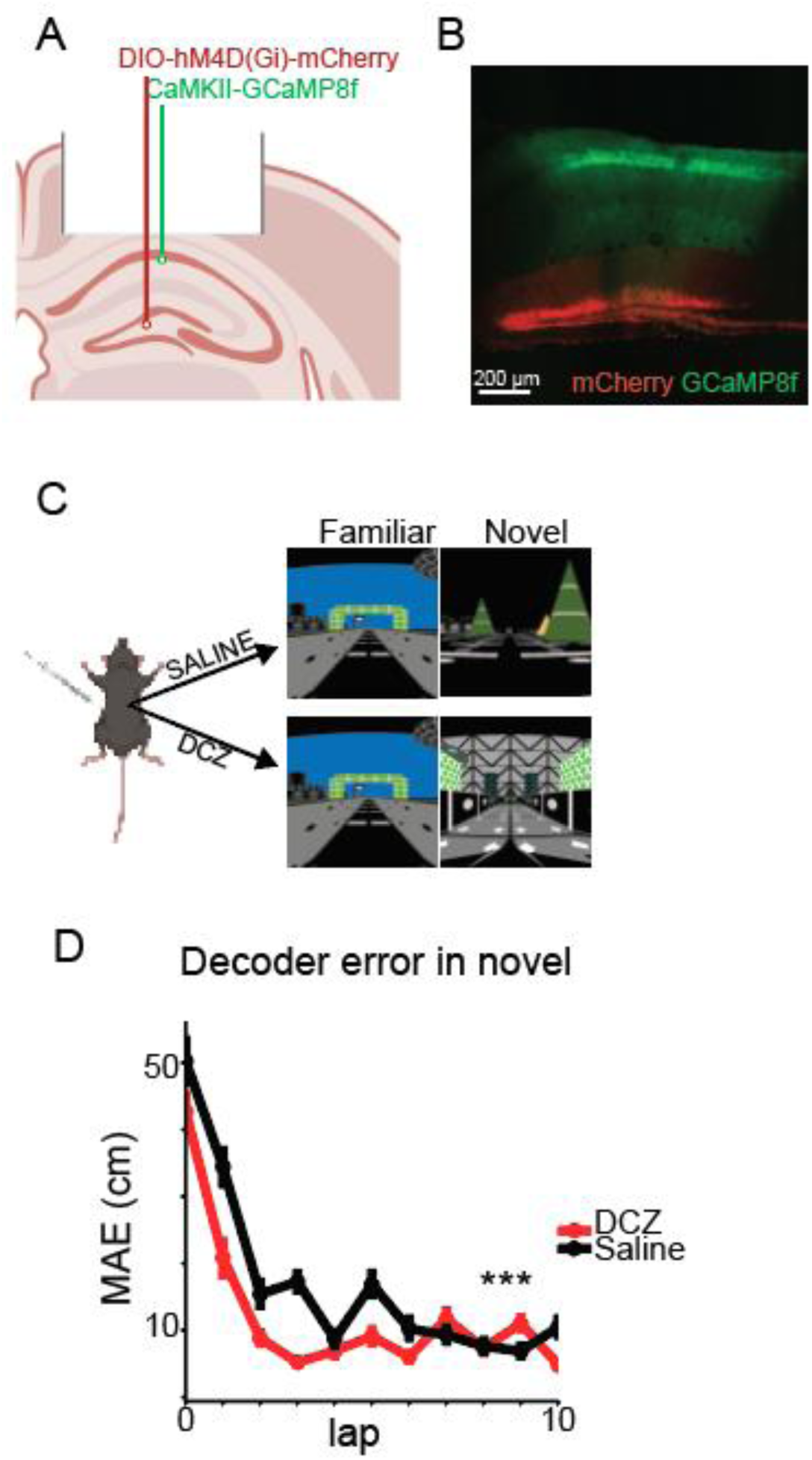
Effects of optogenetic inhibition were not driven by off-target labeling of dentate gyrus granule cells. A-C) protocol for chemogenetic inhibition of granule cells and CA1 imaging in the right hemisphere (adapted from GoodSmith et al., 2025)^57^. A) A cre-dependent inhibitory DREADD virus was injected into the DG, and GCaMP8f was injected into dorsal right CA1. B) mCherry expression can be seen selectively in the granule cell layer and mossy fiber projections to CA1, while GCaMP expression is restricted to CA1. C) On two seperate days, mice (n=4) were injected with DCZ (to inhibit granule cells) or saline (as a within-animal control). On each day, mice were exposed to a novel and familiar environment in ∼5.5 minute blocks. D) Effect of granule cell inhibition on decoding error (similar to Fig 2). In contrast to the increased decoding error observed following optogenetic inhibition of CA3 activity in the early phase, granule cell inhibition significantly decreased decoding error compared to control. (n observations = 1694, ranksum test p <0.001)

**Supp Fig 5.**
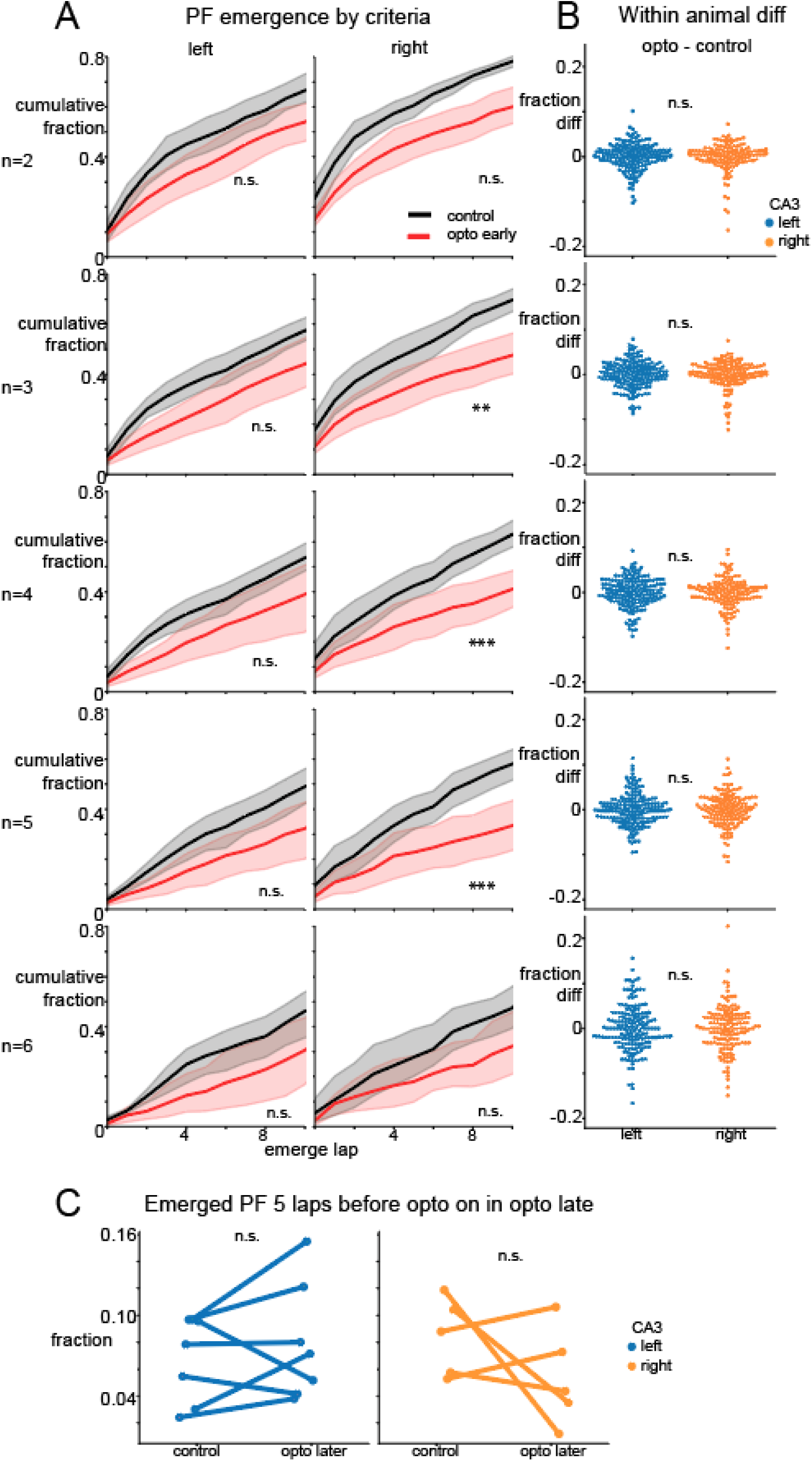
Left and right CA3 input inhibition similarly delayed the emergence of CA1 place cells, regardless of emergence criteria. A. Cumulative histogram of fractions of place fields emerged by lap number comparing opto early (red) with control (black) for first 10 laps under optogenetic inhibition by difference emergence criteria. Emergence lap was defined as the first lap in N out of 6 consecutive laps of within-field firing. P values reported were after Bonferroni correction (*10). N=2, left: Wilcoxon signed-rank stats=701, p =1.2; right: stats= 361.5, p=0.07. N=3, left: stats=612, p=0.26; right: stats=262, p<0.01; N=4, left: stats=588, p=0.16; right: stats=209, p<0.001; N=5, left: stats=543, p=0.062; right: stats=190, p<0.001; N=6, left: stats=506, p=0.11; right: stats=371, p=0.26. B. Within-animal emergence fraction difference between control and opto. Each dot represented one lap: below 0 meant the lap had less fraction of PF emerged under opto than that under lap-matched control, which reflected that optogenetic inhibition delayed the emergence of place fields in that lap. No significant difference found between left and right under any criteria of N. P values reported were after Bonferroni correction (*5). N=2, Mann whitney U stats=1653, p=1.40; N=3, Mann whitney U stats=1702, p=1.23; N=4, T stats = 1.29, p=1; N=5, T stats=1.74, p=0.41; N=6, T stats=0.24, p=4.05. C. Fractions of place fields emerged 5 laps before turning on optogenetic in opto late by animal. No significant differences were found between control and opto in the left or right group. (linear mixed effects model, no significant effects, CA3 p=0.99, opto p=0.46, CA3 x opto p=0.087. Wilcoxon signed-rank test left: p=0.375, n=7, right: p=0.625, n=5.)

**Supp Fig 6.**
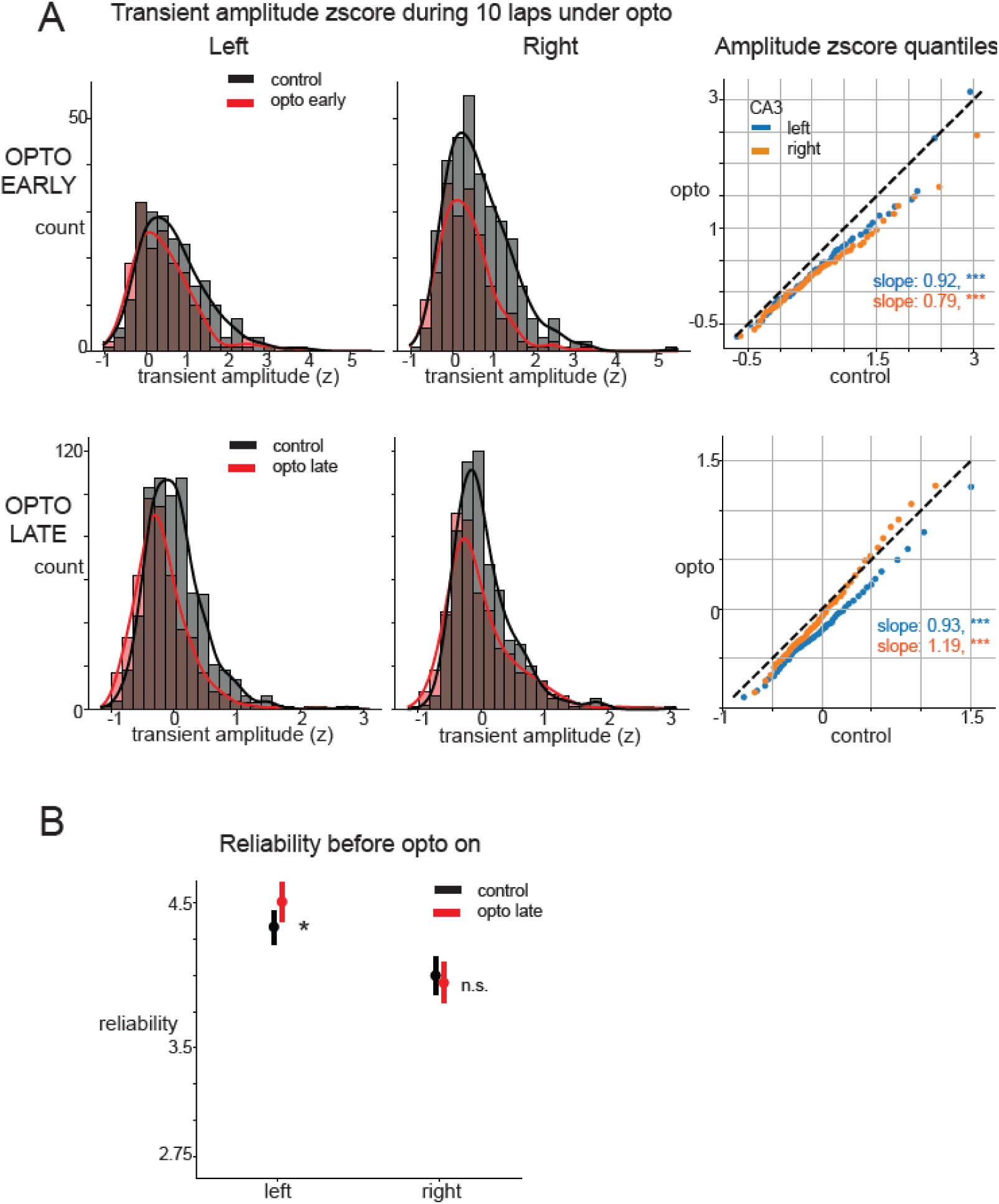
Characterizing place cell amplitude and reliability metrics. A. In-field transient amplitude distribution under opto and control in histogram and qqplot. Top: In opto early, inhibiting left and right CA3 axons during the early phase disproportionately reduced high amplitude transients. Low amplitude transients were not reduced. Left n transients opto=166, control=211; right n transients opto=188, control=332. Quantile-quantile plot compared the distribution of z-scored transient amplitudes in control and opto. A linear regression was fitted to the qqplot. Left: R=0.98, p<0.001, slope=0.92, intercept=-0.17; Right: R=0.99, p<0.001, slope=0.79, intercept=-0.17. Dashed line showed if opto and control had the same distribution for reference, slope smaller than 1 reflects a bigger effect on higher amplitude transients, 1 reflects same effect on all transients regardless of amplitude, bigger than 1 reflects a bigger effect on lower amplitude transients. Bottom: Same as top in opto late. In the later phase, left CA3 input inhibition equally reduced all transients independent of their amplitude. However, right CA3 input inhibition mostly affected low amplitude transients. Left n transients opto=468, control=635; right n transients opto=465, control=603; qqplot. Left: R=0.99, p<0.001, slope=0.93, intercept=-0.2; Right: R=0.99, p<0.001, slope=1.19, intercept=-0.05. B. In-field reliability 5 laps before opto on in opto late condition as control. Right CA3 animals showed no difference from controls. left CA3 animals exhibited slightly higher reliability than controls – opposite to the direction caused by inhibition. Left: N place fields control=781, opto=612, Mann whitney U p=0.034 after Bonferroni correction (*2); Right: N place fields control=818, opto=656, p=0.88.

**Supp Fig 7.**
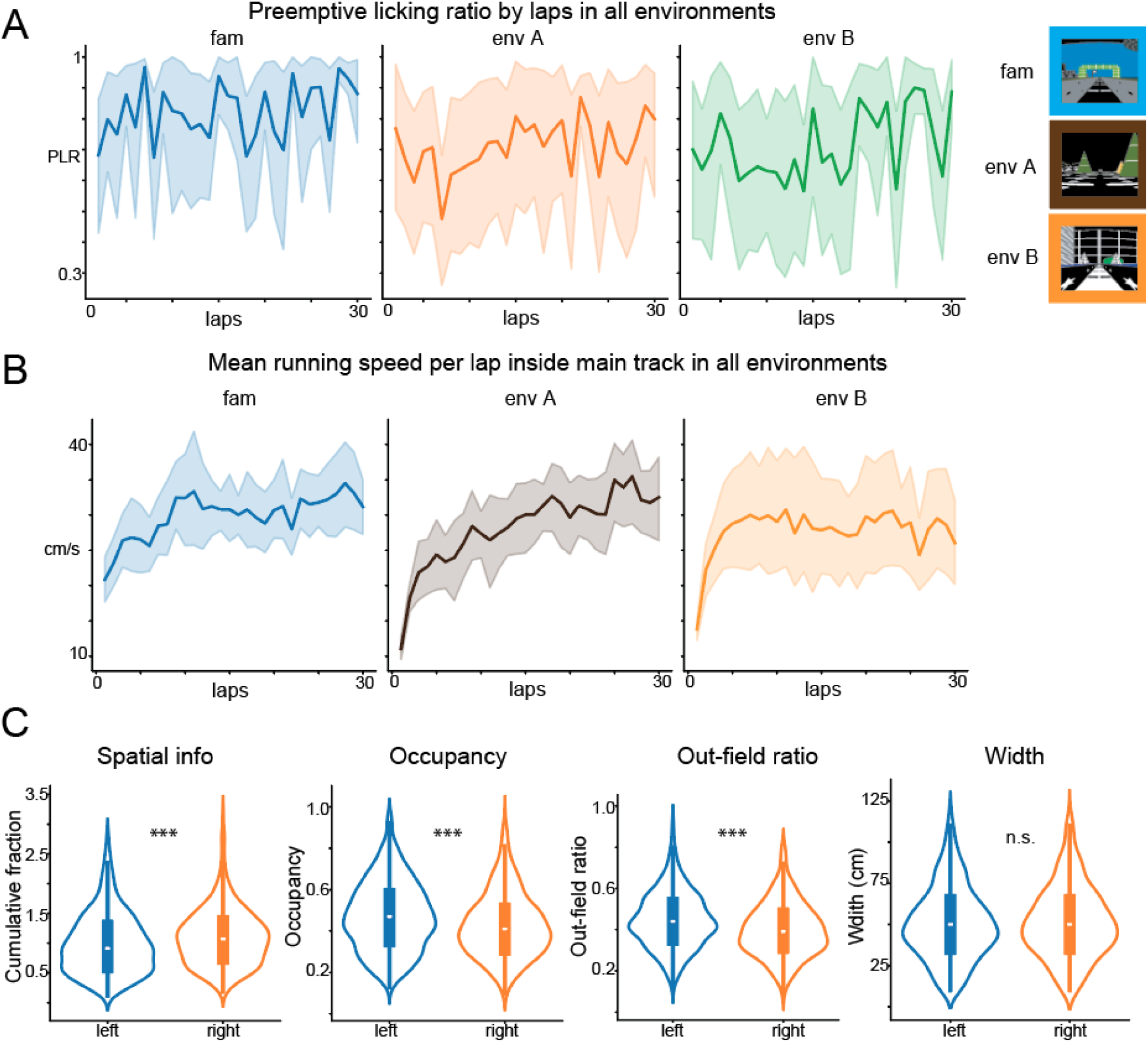
Axon animal behavior and place fields features. A. Mean preemptive licking ratio calculated the same way as Fig1 in axon animals (n=8) in all environments. B. Mean running speed inside the same main track region as Supp. Fig.1 in axon animals (n=8) over laps in all environments. C. Left and right CA3 axon place fields (left n=813, right n=530) feature comparison. Right CA3 place fields contained significantly more spatial information. Mann Whitney U stats=450693, p<0.001. Left CA3 axon place fields had significantly longer occupancy than right CA3 axon place fields. Mann Whitney U stats=607665, p<0.001. Left CA3 axon place fields had significantly more out-field firing ratio than right CA3 axon place fields. Mann Whitney U stats=633514, p<0.001. Left and right CA3 axon place fields were not significantly different from each other. Mann Whitney U stats=514315, p=0.38

**Supp Fig 8.**
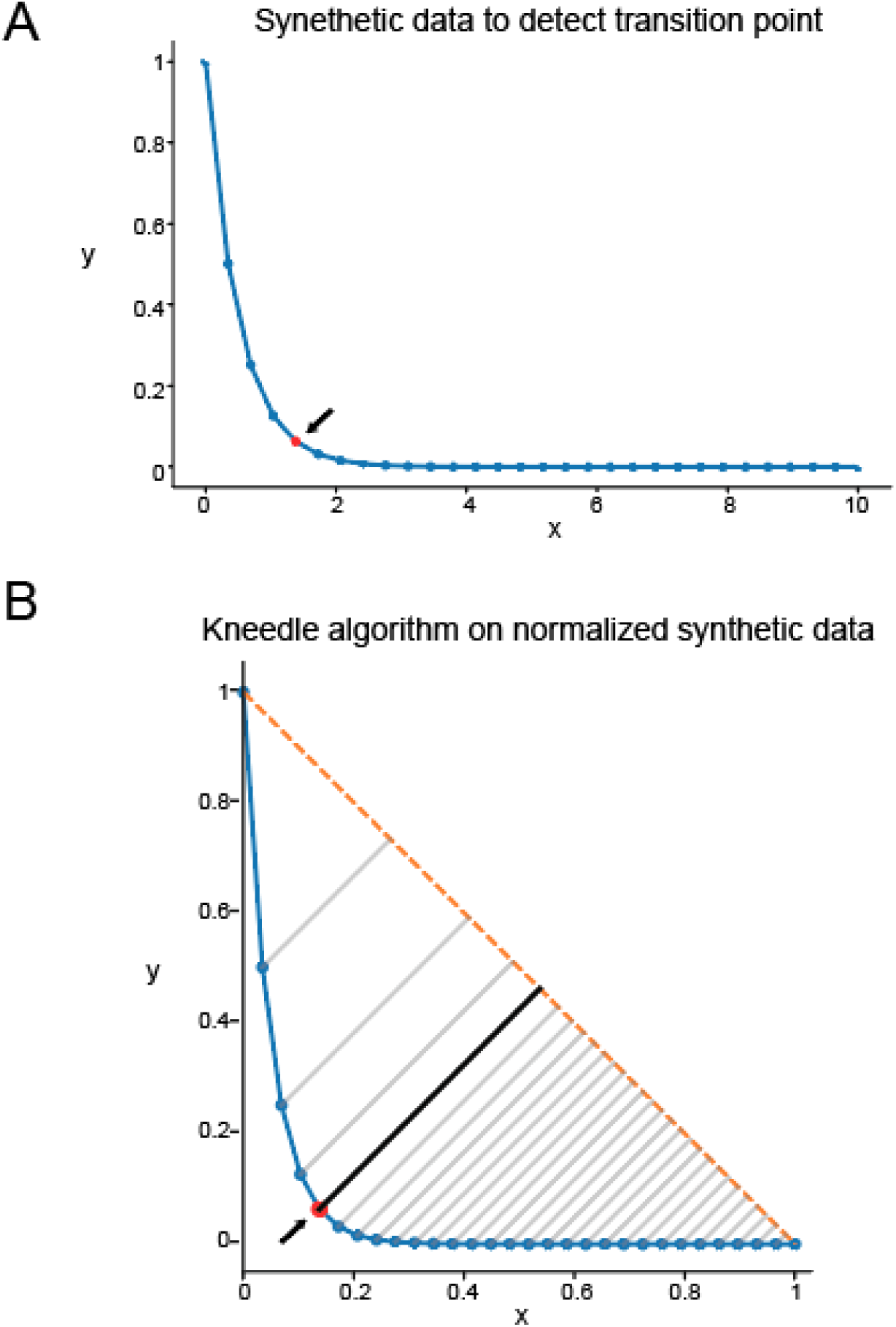
Illustration of kneedle algorithm. A. Example synthetic data with an exponential decay profile used to illustrate detection of a transition point. Arrow pointing at the transition point (red). B. Same dataset after normalization and application of the Kneedle algorithm. The method identifies the point of maximum deviation between the data curve (blue) and the diagonal line (orange dashed) connecting the start and end of the curve (smoothing as needed when using real data). The knee point corresponds to the location with the largest perpendicular distance (black line) from this diagonal, highlighted by the red point and arrow.

## Methods

### Subjects

Some data from the same mice used in this study were analyzed in a previously published paper^16^. Only the broad effect of CA3 input inhibition on CA1 pyramidal firing (reduced somatic activity) was previously reported from this shared dataset. All other analyses and results reported in the current manuscript are new. All experimental and surgical procedures were in accordance with the University of Chicago Animal Care and Use Committee guidelines. For this study, 11-15 week old C57BL/6-Tg(Grik4-cre)G32-4Stl/J mice were individually housed in a reverse 12 hour light/dark cycle in controlled temperature of ∼20 Celsius and ∼50% humidity animal facility. Both male and female mice were included in the experiments. All training and experiments were conducted during the animal’s dark cycle. The entire experimental protocol, from the initial injection to the end of the experiments, would last 4-6 months.

### Injection protocol

Mice were anesthetized and injected with 0.5ml of saline and 0.5ml of meloxicam. A small craniotomy was made over the left or right hippocampus CA3 region (2.0mm caudal, 1.7mm or –1.7mm lateral respectively of Bregma). A Cre-dependent opsin, pAAV-hSyn1-SIO-eOPN3-mScarlet-WPRE (Addgene #125713), was injected into CA3 (∼100nl at a depth of 1.9mm below the dura). Right after the viral injection, the site was covered with dental cement (Metabond, Parkell Corporation) and a metal head plate (Atlas Tool and Die Works). After at least 2 weeks, another small craniotomy was made over the right CA1 region (1.7mm lateral, –2.3mm caudal of Bregma), where a genetically encoded calcium indicator, AAV1-CamKII-GCaMP6f (Addgene #100834) was injected using a beveled glass micropipette. Water restriction (0.8-1ml per day) began after a week of the initial craniotomy. After one week of water restriction, mice went through another surgery to implant a hippocampal cannula window at the right CA1 region. During the surgery, a head plate and a head ring were attached to the cannula window to house the microscope objective and block out ambient light. Post-surgery, water restricted was paused (back to ad lib water access) for one week. Expression of eOPN3 in CA3 terminals and GCaMP6f in pyramidal cells at CA1 was checked every week starting 4 weeks after injection.

For axon imaging, the same injection protocol was followed except for a different virus at a different expression timeline. The genetically-encoded calcium indicator, AAV9-axon-GCaMP6s-P2A-mRuby3 (Addgene #112005) was injected into the left or right CA3 of Grik4-Cre mice (2mm caudal, 1.7mm or –1.7mm lateral and 200 nL at a depth of 1.9 mm below the surface of the dura). After at least 2 weeks post-injection, mice underwent water restriction for a week before the cannula implantation surgery. Expression was checked every week starting 10 weeks after injection.

### Immunofluorescence

Mice were anesthetized with isoflurane and perfused with 15mL of PBS followed by 20mL of 4% PFA. Brains remained in 4% PFA overnight and were then transferred to 30% sucrose. Once brains had sunk in sucrose, they were embedded in OCT and sectioned with a cryostat at a thickness of 30um. A series of every fourth section was collected into a well plate containing PBS.

Free-floating sections were washed in PBS, followed by a 2 hour incubation at room temperature in blocking solution (10% NGS, 0.1% Triton X, 1% BSA). Sections were washed again in PBS and incubated overnight at 4C in 1:250 rabbit anti-PCP4 (PA5-52209, Invitrogen) diluted in blocking solution. A final wash was performed and sections were then incubated for 2 hours at room temperature in secondary antibody 1:500 Alexa Fluor 647 goat anti-rabbit (A21244, Invitrogen) diluted in blocking solution. Sections were collected onto glass slides and mounted with DAPI mounting media (AB104139, Abcam).

Imaging was performed at the University of Chicago Integrated Light Microscopy Core RRID: SCR_019197. Digital image files were created with an Olympus VS200 Research Slide Scanner (Olympus / Evident, Center Valley, PA) with a Hamamatsu ORca-Fusion camera (Hamamatsu Photonics, Skokie, IL). Individual images were created with the OlyVIA Viewer software (Olympus / Evident, Center Valley, PA).

### Histological Analysis

All fluorescence analysis was performed using QuPath version 0.5.1^67^. To quantify eOPN3-mScarlet expression in the DG and CA2 neurons ipsilateral to the injection, individual cells were identified by their presence in the granule cell layer or labeling with the anti-PCP4 antibody, respectively, and the fluorescence intensity was measured. Three sections were used per mouse, and the mean fluorescence intensity was calculated. Intensity was normalized by dividing DG and CA2 fluorescence intensity values by the fluorescence intensity within CA3 somas for each mouse.

### Behavioral training

Animals were trained to run on a treadmill in a virtual reality (VR) environment composed of five LED screens. All VR environments (one training environment, named as familiar environment, and three novel environments env A, B, C) were created using VIRMEn. Each environment had a two-meter linear track with unique detailed distal and proximal 3D visual cues. At the end of the track in all environments, 4 microliter water rewards were delivered and a short VR pause of 1.5s was implemented to allow for water consumption before “teleported” back to the beginning of the track. During training, mice were placed in the familiar environment for ∼30 minutes each day to learn to run and lick the water rewards. Mice had to run at least 2 laps/min for 3 days in a row to proceed to imaging.

### Two-photon imaging and optogenetic inhibition

Imaging was done using a laser scanning two-photon microscope (Neurolabware). The microscope consisted of an 8 KHz resonant scanning module (Thorlabs), a 16×/0.8 NA/3 mm WD water immersion objective (MRP07220, Nikon). GCaMP6f was excited at 920 nm with a femtosecond-pulsed two-photon laser (Insight DS + Dual, Spectra-Physics) and the fluorescence was collected using a GaAsP PMT (H11706, Hamamatsu). Average laser power measured at the objective was ∼80mw each day and kept constant between days of imaging in the same animal. Sampling rate was 31Hz with bidirectional scanning. Opto-stimulation consisted of 625nm light delivered by a LED (Thorlabs) through the microscope objective, with pulse width of 30ms (during which 2P-imaging was stopped), pulse margin of 5.25ms (during which 2P-imaging was performed), frequency of 28.37Hz, and ∼3 minutes of duration. A shutter stopped imaging during an opto-pulse but scanning continued at the same speed. The asynchrony between opto-stimulation and scanning resulted in missing lines for some frames. Each region of interest (ROI) had a slightly different effective sampling rate from the varying number of missing scan lines. The lowest effective sampling rate for a given ROI was ∼10Hz during opto-stimulation. The PicoScope Oscilloscope (PICO4824, Pico Technology) collected the signal from the microscope to synchronize frame acquisition timing with behavior. Mice behavior including treadmill running speed, position, and licking was collected by Picoscope Oscilloscope and synchronized with the imaging for analysis.

For axon imaging, the same setup was used except for the following adjustments: A reduced sampling rate of 15Hz with unidirectional scanning. Average laser power measured at the objective was ∼90mw each day. Axon-GCaMP6s was excited at 920nm and mRuby was excited at 1040nm. Emitted fluorescence was collected using two GaAsP PMTs.

### Imaging sessions

Once mice met the behavioral criteria described above, they underwent test imaging sessions to assess the expression and functionality of eOPN3 and GCaMP6f. eOPN3 expression was confirmed in the basal and apical layers of CA1 using the red anatomical channel. GCaMP6f expression was verified based on the following criteria: absence of resting fluorescence in the nucleus, fast transient kinetics, and normal soma morphology.

During each test imaging session, mice ran for ∼10 minutes in a dark environment (no visual cues and rewards) followed by ∼30 minutes in the familiar environment. Optogenetic inhibition was delivered for 3 minutes once the mouse had been running for at least 3 minutes in either environment. The 6-day imaging protocol on the same field of view was initiated only after confirming a clear reduction in CA1 pyramidal cell activity during optogenetic inhibition in both the dark and familiar environments (Fig. 3d).

During the 6-day imaging protocol, each imaging day consisted of navigation in the familiar environment (8 min) before switching to a novel environment A, B or C (14 min). Each novel environment was paired with an optogenetic inhibition condition: opto-early, opto-late, or control condition (no opto delivered). Each pairing ran for 2 consecutive days. The order and association of novel environments and optogenetic conditions was counterbalanced between animals. On opto-early days, optogenetic inhibition was delivered as soon as the mouse entered the novel environment and lasted for ∼3 minutes. On opto-late days, optogenetic inhibition was delivered for ∼3 minutes after the mouse had completed at least 15 laps in the novel environment. No optogenetic inhibition was delivered in control condition. Imaging was recorded from the same field of view (FOV) throughout the protocol. After each imaging session, multiple averaged FOVs were saved as references for alignment on subsequent days. Data from sessions with noticeable misalignment between days were discarded. For the opto-early condition, one mouse (left CA3 group) was excluded due to noticeable z-drift during the opto early session, but recordings from the same mouse in the opto-late condition were retained.

For axon imaging, mice followed the same behavioral criteria, and axon GCaMP and mRuby expression were confirmed before proceeding. Each day, mice ran in the familiar environment (∼5 minutes) before transitioning to a novel environment (∼10 minutes). To increase axon collection per animal, recordings were performed in two novel environments. Axon activity from the first day exposure in both novel environments was combined for analysis.

### Image processing and ROI selection

Imaging planes acquired on different days were combined and preprocessed using Suite2p ^53^. Time-series images went through two times of rigid and non-rigid transformations in Suite2p to remove movement artifacts. Motion corrected videos were visually assessed to ensure the absence of drifts in z-direction. Datasets with visible z-drifts were discarded. ROIs were identified by Suite2p and manually inspected for accuracy. For each ROI, baseline corrected ΔF/F traces were generated within each day, filtered for significant calcium transients as reported before. When generating baseline corrected ΔF/F traces for a given ROI, missing frames in optogenetic conditions were linearly interpolated using the *fillmissing* function in MATLAB. To match optogenetic conditions, two control datasets were generated: one downsampled at opto-early inhibition periods and the other at opto-late inhibition periods.

For axon imaging, ROIs were first defined using Suite2p and then preprocessed using default parameters in package *subprep*^54^. A red mRuby channel was recorded simultaneously to axon GCaMP for motion correction and motion artifacts removal.

### Granule cell inhibition experiments

To control for the effects of possible Grik-4 expression in a subset of dentate gyrus granule cells, we compared our results to the effects of selective granule cell inhibition. The data for this analysis was collected as part of a different study (cite) that used a chemogenetic approach to inhibit granule cell activity while recording CA1 place cell activity. Four mice from this study completed enough laps across all conditions to be included in our analysis. To selectively inhibit granule cells, a cre-dependent inhibitory DREADD virus (hSyn-DIO-hM4D(Gi)-mCherry; Addgene, #44362) was injected into the right dorsal dentate gyrus of Dock10-cre mice^66^ (gift from Tonegawa lab). In the same surgery, GCaMP8f (CAMKIIa-jGCaMP8f; Addgene, #176750) was injected in the dorsal CA1 at the same coordinates. Following cannula implantation and recovery, mice were trained to run for water rewards as described above.

On two separate imaging days, mice either received an IP injection of 0.1mg/kg Deschloroclozapine dihydrochlorine (DCZ; to inhibit granule cells) or an equal volume of saline (as a within-animal control). Each imaging day consisted of a ∼5.5 minute session in a familiar environment (the same environment used for training), followed by a ∼5.5 minute session in a novel environment (different novel environment on each day). Place cells were identified using the method described in GoodSmith et al (2025)^57^. Briefly, any cell with a statistically significant spatial information score and a defined place field onset were identified as place cells.

### Transient Raster and Normalized Transient Frequency

To ensure behavioral consistency, neural data during stationary periods was excluded. Control data was downsampled during the same frames as opto-on periods in opto-early and opto-late conditions, ensuring a matched sampling rate during optogenetic inhibition. The data was then binned into ∼160 ms time bins and z-score transformed (described as “z-score transformed neural data” in the study). Time bins with a z-score > 1 were classified as active and counted toward the number of active transients. Transient frequency for each pyramidal cell was calculated as the number of active transients divided by the condition duration. Normalized transient frequency was computed as the transient frequency during opto-on (or time-matched control) divided by the transient frequency of the entire session.

### Low-Dimensional Manifold Visualization with CEBRA

CEBRA was applied on the z-score transformed neural data using environment labels and track location as behavioral variables, with each point in the low-dimensional manifold representing population activity within a single time bin.

To compute embedding centroids for each environment, the 2 m track was divided into 40 spatial bins (5 cm each). Within each bin, the centroid was defined as the average position of all points from the same lap in the latent space. Distance to centroid was calculated as the Euclidean distance between each point and its corresponding centroid based on track location. To ensure equal representation of time bins across conditions, opto-off time bins were randomly downsampled to match the number of opto-on time bins. The average distance to centroid within each opto condition was computed after 20 shuffles.

### Spatial Location Decoding with CEBRA

Similar to manifold visualization, z-score transformed neural data was used. In each shuffle, 80% of the opto-off period was randomly selected to generate the low-dimensional embedding, which was used to train a k-nearest neighbor (KNN) regressor. The model was then tested on 80% of the opto-on period projected embedding. Each animal underwent 20 shuffles, and hyperparameters for both the KNN regressor and latent embedding were optimized via grid search. Within-animal controls were time-matched to the opto-on period in both opto-early and opto-late conditions. Decoding accuracy at each time bin was measured using mean absolute error (MAE), calculated as the absolute difference (in cm) between predicted and true locations. To examine the temporal evolution of spatial decoding accuracy, mean MAE was averaged across all time bins within each lap. To assess the impact of optogenetic inhibition on decoder performance, the within-animal difference in mean MAE between opto and time-matched control conditions was computed for each shuffle (Fig. 4d).

### Spatial Location Decoding with Bayesian Decoder

The 200 cm track was divided into 40 spatial bins (5 cm each). Place cell activity was binarized based on spatial bins and used to train and test a Gaussian Naive Bayes decoder. A cross-validation strategy was applied over 500 shuffles, with 80% of laps randomly selected for training and the remaining 20% for testing in each shuffle. Decoding error was calculated as the absolute difference between predicted and true location bins (e.g., a 2-bin difference corresponds to a 10 cm error). Within each animal, mean decoder error was averaged across all locations within the same lap over all shuffles.

### Licking Behavior

Licking data was recorded using a capacitive sensor on the waterspout. In familiar environments, well-trained mice exhibited increased licking in the pre-reward zone, the region just before reward delivery (Fig. 1). To identify this zone, we averaged lick counts by location across all animals in the familiar environment and applied the kneedle algorithm to detect the transition to elevated anticipatory licking. The pre-reward zone extended from this transition point to just before reward delivery and was consistently applied across all animals and environments. Similarly, the main track zone was determined using the kneedle algorithm to identify the shift from elevated consummatory licking to stable no licking in the familiar environment. Preemptive licking ratio is a lap-by-lap measurement of the ratio between lick count in the pre-reward zone and main track zone.

### Kneedle Algorithm to Determine Novelty Modulation Time

Implemented and adapted from the Kneedle algorithm ^31^, this procedure was applied to each time series (e.g. Bayesian decoder error by lap, transient raster count over time etc). Following preprocessing (smoothing as described and normalized), a linear trend was fitted between the first and last time points of the signal. For each intermediate time point, the perpendicular distance to this line was computed. The time point with the maximum deviation—i.e., the “elbow” point—was defined as the novelty modulation time, marking the transition from a phase of rapid decline to a stable period. See Supp Fig 8 for visualization.

### Place Field Selection

We used the peak method ^54^ with additional criteria to account for noise in the dataset. Potential place fields (PFs) were identified as contiguous regions on the mean tuning curve with a minimum amplitude of 0.1 ΔF/F and exceeding 40% of the difference between peak ΔF/F and baseline values. At least 30% of laps had to contain a significant transient within the PF boundaries, and PF width was constrained to 6%–50% of the track length. Each PF had to be active within-field in at least 3 of 6 consecutive laps (see emergence lap below), but only one instance of this criterion being met was required, defined as the emergence lap. After passing all those criteria, neural activity was randomly shuffled 600 times by shifting fluorescence transients in time while preserving their calcium dynamics. For each shuffle, peak and bottom fluorescence values were averaged across the fluorescence map. Cells with true peak fluorescence in the top 1% and bottom fluorescence in the bottom 1% were classified as place cells.

### Emergence lap

For each cell, the first lap in n (n=3 if not otherwise specified) out of 6 consecutive laps of within-field firing is defined as an emergence lap.

### Reliability

For each place cell, reliability is defined as the number of laps with significant in-field transient over a 6-lap sliding window.

### Transient Amplitude

Neural activity traces were smoothed using a centered rolling average with a ∼320 ms time window (minimum ∼100 ms per window) to reduce high-frequency noise. Peaks were then detected using *scipy.find_peaks* with the following criteria: Minimum prominence of 0.1ΔF/F, minimum peak height of 0.12ΔF/F, minimum interpeak interval of ∼320ms, peak width of ∼160 ms to ∼3.88s. Transient amplitude was defined as the area under the curve from transient onset to peak.

### Normalized axon activity

Calcium transient amplitudes were calculated as above but with lower prominence (0.05 ΔF/F) and minimum peak height (0.1 ΔF/F). Normalized axon activity was defined as the sum of all transient amplitudes divided by the number of laps (10 laps in early, 15 laps in late), capturing both transient frequency and amplitude.

### Statistics and Reproducibility

Error bars in all figures represent 95% confidence intervals of a bootstrap-generated difference (5,000 resamples). Data distributions were assessed for normality using the Shapiro–Wilk test. If normally distributed, a paired or unpaired Student’s t-test was used where applicable. For non-normal distributions, a paired Wilcoxon signed-rank test or an unpaired Mann–Whitney U-test was applied. For samples with six or fewer data points, only non-parametric tests were used. Multiple comparisons were corrected using Bonferroni post hoc adjustments. p < 0.05 was chosen to indicate statistical significance. P values presented in figures are as follows: *p < 0.05, **p < 0.01, ***p < 0.001, n.s. not significant.

To compare delta-delta differences (Fig. 4d, 5b inset, 5c inset, 5h inset, 5i inset, 5k, 5l) and assess whether left or right inhibition had a greater effect, a within-animal subsampled bootstrap difference distribution was generated. For each shuffle, an equal number of subsamples were randomly selected from the control and opto conditions within the same animal, and their mean difference was computed. This process was repeated 500 times per animal to construct a bootstrap distribution of control–opto differences. Within-animal differences from the same inhibition group were pooled for left vs. right comparisons, and statistical significance was tested accordingly.

A linear mixed effects model (Fig. 5E bottom, 5F bottom, Supp. Fig. 3c) was implemented in MATLAB using *fitlme* with the formula: *EmergePercent ∼ CA3 ∗ opto + (1|mouse)*.

This model assesses the effects of optogenetic manipulation, CA3 side, and their interaction while accounting for variability between animals.

## Acknowledgements

We thank Leslie Kay, PhD and Seetha Krishnan, PhD for feedback on the manuscript and all members of the Sheffield lab for valuable discussions that helped shape the manuscript, especially Roma Shah for assistance in animal training and Mikayla Voglewede, PhD for histology analysis.

## Author contributions

A.J. and M.S. conceived and designed the experiments. A.J. collected the optogenetic and axon data. A.J. analyzed the data. J.R.M. and A.T. collected the histology and immunofluorescence data. J.R.M. analyzed the histology and immunofluorescence data. D.G. collected and analyzed the dentate gyrus data. A.J. and M.S. interpreted the data and wrote the original manuscript. A.J., D.G., J.R.M., A.T., M.S. reviewed and edited the manuscript. M.S. supervised the research and obtained funding.

## Data and code availability

The original codes have been deposited at https://github.com/anqijiang/opto_analysis and will be publicly available on the date of publication. Raw imaging data is large and not feasible for upload to an online repository but is available upon request to the lead contact at. Processed source data for all figures and associated statistical analysis will be uploaded to Box and provided in the final version of the paper. Any additional information required to reanalyze the data reported in this paper is available from the lead contact upon request.

## References

1. Eichenbaum, H., Dudchenko, P., Wood, E., Shapiro, M., and Tanila, H. (1999). The Hippocampus, Memory, and Place Cells Is It Spatial Memory or a Memory Space? Neuron 23, 209–226. 10.1016/s0896-6273(00)80773-4.

2. Robinson, N.T.M., Descamps, L.A.L., Russell, L.E., Buchholz, M.O., Bicknell, B.A., Antonov, G.K., Lau, J.Y.N., Nutbrown, R., Schmidt-Hieber, C., and Häusser, M. (2020). Targeted Activation of Hippocampal Place Cells Drives Memory-Guided Spatial Behavior. Cell 183, 1586–1599.e10. 10.1016/j.cell.2020.09.061.

3. O’Keefe, J., and Nadel, L. (1979). Précis of O’Keefe & Nadel’s The hippocampus as a cognitive map. Behav. Brain Sci. 2, 487–494. 10.1017/s0140525x00063949.

4. Leussis, M.P., and Bolivar, V.J. (2006). Habituation in rodents: A review of behavior, neurobiology, and genetics. Neurosci. Biobehav. Rev. 30, 1045–1064. 10.1016/j.neubiorev.2006.03.006.

5. Wilkinson, J.L., Herrman, L., Palmatier, M.I., and Bevins, R.A. (2006). Rats’ novel object interaction as a measure of environmental familiarity. Learn. Motiv. 37, 131–148. 10.1016/j.lmot.2005.04.001.

6. Bevins, R.A., Koznarova, J., and Armiger, T.J. (2001). Environmental Familiarization in Rats: Differential Effects of Acute and Chronic Nicotine. Neurobiol. Learn. Mem. 75, 63–76. 10.1006/nlme.1999.3955.

7. Dong, C., Madar, A.D., and Sheffield, M.E.J. (2021). Distinct place cell dynamics in CA1 and CA3 encode experience in new environments. Nat. Commun. 12, 2977. 10.1038/s41467-021-23260-3.

8. Berners-Lee, A., Feng, T., Silva, D., Wu, X., Ambrose, E.R., Pfeiffer, B.E., and Foster, D.J. (2022). Hippocampal replays appear after a single experience and incorporate greater detail with more experience. Neuron 110, 1829–1842.e5. 10.1016/j.neuron.2022.03.010.

9. Cohen, J.D., Bolstad, M., and Lee, A.K. (2017). Experience-dependent shaping of hippocampal CA1 intracellular activity in novel and familiar environments. eLife 6, e23040. 10.7554/elife.23040.

10. Chiu, Y., Dong, C., Krishnan, S., and Sheffield, M.E.J. (2023). The Precision of Place Fields Governs Their Fate across Epochs of Experience. eNeuro 10, ENEURO.0261-23.2023. 10.1523/eneuro.0261-23.2023.

11. Zemla, R., Moore, J.J., Hopkins, M.D., and Basu, J. (2022). Task-selective place cells show behaviorally driven dynamics during learning and stability during memory recall. Cell Rep. 41, 111700. 10.1016/j.celrep.2022.111700.

12. Sheffield, M.E.J., Adoff, M.D., and Dombeck, D.A. (2017). Increased Prevalence of Calcium Transients across the Dendritic Arbor during Place Field Formation. Neuron 96, 490–504.e5. 10.1016/j.neuron.2017.09.029.

13. Geiller, T., Sadeh, S., Rolotti, S.V., Blockus, H., Vancura, B., Negrean, A., Murray, A.J., Rózsa, B., Polleux, F., Clopath, C., et al. (2022). Local circuit amplification of spatial selectivity in the hippocampus. Nature 601, 105–109. 10.1038/s41586-021-04169-9.

14. Bittner, K.C., Milstein, A.D., Grienberger, C., Romani, S., and Magee, J.C. (2017). Behavioral time scale synaptic plasticity underlies CA1 place fields. Science 357, 1033–1036. 10.1126/science.aan3846.

15. Bittner, K.C., Grienberger, C., Vaidya, S.P., Milstein, A.D., Macklin, J.J., Suh, J., Tonegawa, S., and Magee, J.C. (2015). Conjunctive input processing drives feature selectivity in hippocampal CA1 neurons. Nat. Neurosci. 18, 1133–1142. 10.1038/nn.4062.

16. Madar, A.D., Jiang, A., Dong, C., and Sheffield, M.E.J. (2025). Synaptic plasticity rules driving representational shifting in the hippocampus. Nat. Neurosci., 1–13. 10.1038/s41593-025-01894-6.55.

17. Priestley, J.B., Bowler, J.C., Rolotti, S.V., Fusi, S., and Losonczy, A. (2022). Signatures of rapid plasticity in hippocampal CA1 representations during novel experiences. Neuron 110, 1978–1992.e6. 10.1016/j.neuron.2022.03.026.

18. Kesner, R.P. (2007). Behavioral functions of the CA3 subregion of the hippocampus. Learn. Mem. 14, 771–781. 10.1101/lm.688207.

19. Nakazawa, K., Quirk, M.C., Chitwood, R.A., Watanabe, M., Yeckel, M.F., Sun, L.D., Kato, A., Carr, C.A., Johnston, D., Wilson, M.A., et al. (2002). Requirement for Hippocampal CA3 NMDA Receptors in Associative Memory Recall. Science 297, 211–218. 10.1126/science.1071795.

20. Li, X.-G., Somogyi, P., Ylinen, A., and Buzsáki, G. (1994). The hippocampal CA3 network: An in vivo intracellular labeling study. J. Comp. Neurol. 339, 181–208. 10.1002/cne.903390204.

21. Kohl, M.M., Shipton, O.A., Deacon, R.M., Rawlins, J.N.P., Deisseroth, K., and Paulsen, O. (2011). Hemisphere-specific optogenetic stimulation reveals left-right asymmetry of hippocampal plasticity. Nat. Neurosci. 14, 1413–1415. 10.1038/nn.2915.

22. Shinohara, Y., Hirase, H., Watanabe, M., Itakura, M., Takahashi, M., and Shigemoto, R. (2008). Left-right asymmetry of the hippocampal synapses with differential subunit allocation of glutamate receptors. Proc. Natl. Acad. Sci. 105, 19498–19503. 10.1073/pnas.0807461105.

23. Shipton, O.A., El-Gaby, M., Apergis-Schoute, J., Deisseroth, K., Bannerman, D.M., Paulsen, O., and Kohl, M.M. (2014). Left–right dissociation of hippocampal memory processes in mice. Proc. Natl. Acad. Sci. 111, 15238–15243. 10.1073/pnas.1405648111.

24. Guan, H., Middleton, S.J., Inoue, T., and McHugh, T.J. (2021). Lateralization of CA1 assemblies in the absence of CA3 input. Nat. Commun. 12, 6114. 10.1038/s41467-021-26389-3.

25. El-Gaby, M., Zhang, Y., Wolf, K., Schwiening, C.J., Paulsen, O., and Shipton, O.A. (2016). Archaerhodopsin Selectively and Reversibly Silences Synaptic Transmission through Altered pH. Cell Rep. 16, 2259–2268. 10.1016/j.celrep.2016.07.057.

26. Song, D., Wang, D., Yang, Q., Yan, T., Wang, Z., Yan, Y., Zhao, J., Xie, Z., Liu, Y., Ke, Z., et al. (2020). The lateralization of left hippocampal CA3 during the retrieval of spatial working memory. Nat. Commun. 11, 2901. 10.1038/s41467-020-16698-4.

27. El-Gaby, M., Reeve, H.M., Lopes-dos-Santos, V., Campo-Urriza, N., Perestenko, P.V., Morley, A., Strickland, L.A.M., Lukács, I.P., Paulsen, O., and Dupret, D. (2021). An emergent neural coactivity code for dynamic memory. Nat. Neurosci. 24, 694–704. 10.1038/s41593-021-00820-w.

28. Kawakami, R., Shinohara, Y., Kato, Y., Sugiyama, H., Shigemoto, R., and Ito, I. (2003). Asymmetrical Allocation of NMDA Receptor ε2 Subunits in Hippocampal Circuitry. Science 300, 990–994. 10.1126/science.1082609.

29. Jordan, J.T. (2020). The rodent hippocampus as a bilateral structure: A review of hemispheric lateralization. Hippocampus 30, 278–292. 10.1002/hipo.23188.

30. Mahn, M., Saraf-Sinik, I., Patil, P., Pulin, M., Bitton, E., Karalis, N., Bruentgens, F., Palgi, S., Gat, A., Dine, J., et al. (2021). Efficient optogenetic silencing of neurotransmitter release with a mosquito rhodopsin. Neuron 109, 1621–1635.e8. 10.1016/j.neuron.2021.03.013.

31. Satopää, V., Albrecht, J., Irwin, D., and Raghavan, B. (2011). Finding a “Kneedle” in a Haystack: Detecting Knee Points in System Behavior. 2011 31st Int. Conf. Distrib. Comput. Syst. Work., 166–171. 10.1109/icdcsw.2011.20.

32. Davoudi, H., and Foster, D.J. (2019). Acute silencing of hippocampal CA3 reveals a dominant role in place field responses. Nat. Neurosci. 22, 337–342. 10.1038/s41593-018-0321-z.

33. Zutshi, I., Valero, M., Fernández-Ruiz, A., and Buzsáki, G. (2022). Extrinsic control and intrinsic computation in the hippocampal CA1 circuit. Neuron 110, 658–673.e5. 10.1016/j.neuron.2021.11.015.

34. Guo, W., Zhang, J.J., Newman, J.P., and Wilson, M.A. (2024). Latent learning drives sleep-dependent plasticity in distinct CA1 subpopulations. Cell Rep. 43, 115028. 10.1016/j.celrep.2024.115028.

35. Yang, W., Sun, C., Huszár, R., Hainmueller, T., Kiselev, K., and Buzsáki, G. (2024). Selection of experience for memory by hippocampal sharp wave ripples. Science 383, 1478– 1483. 10.1126/science.adk8261.

36. Nieh, E.H., Schottdorf, M., Freeman, N.W., Low, R.J., Lewallen, S., Koay, S.A., Pinto, L., Gauthier, J.L., Brody, C.D., and Tank, D.W. (2021). Geometry of abstract learned knowledge in the hippocampus. Nature 595, 80–84. 10.1038/s41586-021-03652-7.

37. Jones, E.A.A., and Giocomo, L.M. (2023). Neural ensembles in navigation: From single cells to population codes. Curr. Opin. Neurobiol. 78, 102665. 10.1016/j.conb.2022.102665.

38. Schneider, S., Lee, J.H., and Mathis, M.W. (2023). Learnable latent embeddings for joint behavioural and neural analysis. Nature 617, 360–368. 10.1038/s41586-023-06031-6.

39. McKenzie, S., Huszár, R., English, D.F., Kim, K., Christensen, F., Yoon, E., and Buzsáki, G. (2021). Preexisting hippocampal network dynamics constrain optogenetically induced place fields. Neuron 109, 1040–1054.e7. 10.1016/j.neuron.2021.01.011.

40. Keinath, A.T., Nieto-Posadas, A., Robinson, J.C., and Brandon, M.P. (2020). DG–CA3 circuitry mediates hippocampal representations of latent information. Nat. Commun. 11, 3026. 10.1038/s41467-020-16825-1.

41. Ji, D., and Wilson, M.A. (2008). Firing Rate Dynamics in the Hippocampus Induced by Trajectory Learning. J. Neurosci. 28, 4679–4689. 10.1523/jneurosci.4597-07.2008.

42. Kitamura, T., Ogawa, S.K., Roy, D.S., Okuyama, T., Morrissey, M.D., Smith, L.M., Redondo, R.L., and Tonegawa, S. (2017). Engrams and circuits crucial for systems consolidation of a memory. Science 356, 73–78. 10.1126/science.aam6808.

43. Butola, T., Hernández-Frausto, M., Blankvoort, S., Flatset, M.S., Peng, L., Hairston, A., Johnson, C.D., Elmaleh, M., Amilcar, A., Hussain, F., et al. (2025). Hippocampus shapes entorhinal cortical output through a direct feedback circuit. Nat. Neurosci., 1–12. 10.1038/s41593-025-01883-9.

44. Jiang, A., Zhao, C., and Sheffield, M. (2024). A preprocessing toolbox for 2-photon subcellular calcium imaging. bioRxiv, 2024.10.04.616737. 10.1101/2024.10.04.616737.

45. Rebola, N., Carta, M., and Mulle, C. (2017). Operation and plasticity of hippocampal CA3 circuits: implications for memory encoding. Nat. Rev. Neurosci. 18, 208–220. 10.1038/nrn.2017.10.

46. Zaki, Y., and Cai, D.J. (2025). Memory engram stability and flexibility. Neuropsychopharmacology 50, 285–293. 10.1038/s41386-024-01979-z.

47. Geva, N., Deitch, D., Rubin, A., and Ziv, Y. (2023). Time and experience differentially affect distinct aspects of hippocampal representational drift. Neuron 111, 2357–2366.e5. 10.1016/j.neuron.2023.05.005.

48. Keinath, A.T., Mosser, C.-A., and Brandon, M.P. (2022). The representation of context in mouse hippocampus is preserved despite neural drift. Nat. Commun. 13, 2415. 10.1038/s41467-022-30198-7.

49. Khatib, D., Ratzon, A., Sellevoll, M., Barak, O., Morris, G., and Derdikman, D. (2023). Active experience, not time, determines within-day representational drift in dorsal CA1. Neuron 111, 2348–2356.e5. 10.1016/j.neuron.2023.05.014.

50. Krishnan, S., and Sheffield, M.E.J. (2023). Reward Expectation Reduces Representational Drift in the Hippocampus. bioRxiv, 2023.12.21.572809. 10.1101/2023.12.21.572809.

51. Delamare, G., Zaki, Y., Cai, D.J., and Clopath, C. (2023). Drift of neural ensembles driven by slow fluctuations of intrinsic excitability. 10.7554/elife.88053.2.

52. Krishnan, S., Heer, C., Cherian, C., and Sheffield, M.E.J. (2022). Reward expectation extinction restructures and degrades CA1 spatial maps through loss of a dopaminergic reward proximity signal. Nat. Commun. 13, 6662. 10.1038/s41467-022-34465-5.

53. Pachitariu, M., Stringer, C., Dipoppa, M., Schröder, S., Rossi, L.F., Dalgleish, H., Carandini, M., and Harris, K.D. (2017). Suite2p: beyond 10,000 neurons with standard two-photon microscopy. bioRxiv, 061507. 10.1101/061507.

54. Grijseels, D.M., Shaw, K., Barry, C., and Hall, C.N. (2021). Choice of method of place cell classification determines the population of cells identified. PLoS Comput. Biol. 17, e1008835. 10.1371/journal.pcbi.1008835.

55. Mahn, M., Saraf-Sinik, I., Patil, P., Pulin, M., Bitton, E., Karalis, N., Bruentgens, F., Palgi, S., Gat, A., Dine, J., et al. (2021). Efficient optogenetic silencing of neurotransmitter release with a mosquito rhodopsin. Neuron 109, 1621–1635.e8. 10.1016/j.neuron.2021.03.013.

56. Heer, C., and Sheffield, M. (2024). Distinct catecholaminergic pathways projecting to hippocampal CA1 transmit contrasting signals during navigation in familiar and novel environments. eLife 13, RP95213. 10.7554/elife.95213.56.

57. GoodSmith, D., Carson, W.H., and Sheffield, M.E.J. (2025). Complementary regulation of memory flexibility and stabilization by dentate gyrus granule cells and mossy cells. BioRxiv. 10.1101/2025.09.05.674488.

58. Sumegi, M., Olah, G., Lukacs, I.P., Blazsek, M., Makara, J.K., and Nusser, Z. (2025). Diverse calcium dynamics underlie place field formation in hippocampal CA1 pyramidal cells. eLife. 10.7554/elife.103676.2.

59. Krishnan, S., Dong, C., Ratigan, H., Morales-Rodriguez, D., Cherian, C., and Sheffield, M. (2025). A contextual fear conditioning paradigm in head-fixed mice exploring virtual reality. eLife. 10.7554/elife.105422.3.

60. Krishnan, S., Heer, C., Cherian, C., and Sheffield, M.E.J. (2022). Reward expectation extinction restructures and degrades CA1 spatial maps through loss of a dopaminergic reward proximity signal. Nat. Commun. 13, 6662. 10.1038/s41467-022-34465-5.

61. El-Gaby, M., Reeve, H.M., Lopes-dos-Santos, V., Campo-Urriza, N., Perestenko, P.V., Morley, A., Strickland, L.A.M., Lukács, I.P., Paulsen, O., and Dupret, D. (2021). An emergent neural coactivity code for dynamic memory. Nat. Neurosci. 24, 694–704. 10.1038/s41593-021-00820-w.

62. Shipton, O.A., El-Gaby, M., Apergis-Schoute, J., Deisseroth, K., Bannerman, D.M., Paulsen, O., and Kohl, M.M. (2014). Left–right dissociation of hippocampal memory processes in mice. Proc. Natl. Acad. Sci. 111, 15238–15243. 10.1073/pnas.1405648111.

63. Klur, S., Muller, C., Vasconcelos, A.P. de, Ballard, T., Lopez, J., Galani, R., Certa, U., and Cassel, J.-C. (2009). Hippocampal-dependent spatial memory functions might be lateralized in rats: An approach combining gene expression profiling and reversible inactivation. Hippocampus. 10.1002/hipo.20562.

64. El-Gaby, M., Zhang, Y., Wolf, K., Schwiening, C.J., Paulsen, O., and Shipton, O.A. (2016). Archaerhodopsin Selectively and Reversibly Silences Synaptic Transmission through Altered pH. Cell Reports. 10.1016/j.celrep.2016.07.057

65. Li, S., Cullen, W.K., Anwyl, R., and Rowan, M.J. (2003). Dopamine-dependent facilitation of LTP induction in hippocampal CA1 by exposure to spatial novelty. Nature Neuroscience. 10.1038/nn1049.

66. Kohara, K., Pignatelli, M., Rivest, A.J., Jung, H.-Y., Kitamura, T., Suh, J., Frank, D., Kajikawa, K., Mise, N., Obata, Y., et al. (2014). Cell type–specific genetic and optogenetic tools reveal hippocampal CA2 circuits. Nature Neuroscience. 10.1038/nn.3614.

67. Bankhead, P., Loughrey, M.B., Fernández, J.A., Dombrowski, Y., McArt, D.G., Dunne, P.D., McQuaid, S., Gray, R.T., Murray, L.J., Coleman, H.G., et al. (2017). QuPath: Open source software for digital pathology image analysis. Scientific Reports. 10.1038/s41598-017-17204-5.

68. Kitanishi, T., Ujita, S., Fallahnezhad, M., Kitanishi, N., Ikegaya, Y., and Tashiro, A. (2015). Novelty-Induced Phase-Locked Firing to Slow Gamma Oscillations in the Hippocampus: Requirement of Synaptic Plasticity. Neuron. 10.1016/j.neuron.2015.05.012.

