## Supplementary information for "Learning-Dependent Shift from Right to Left CA3 Input Dominance Shapes the Evolution of Right CA1 Spatial Maps"

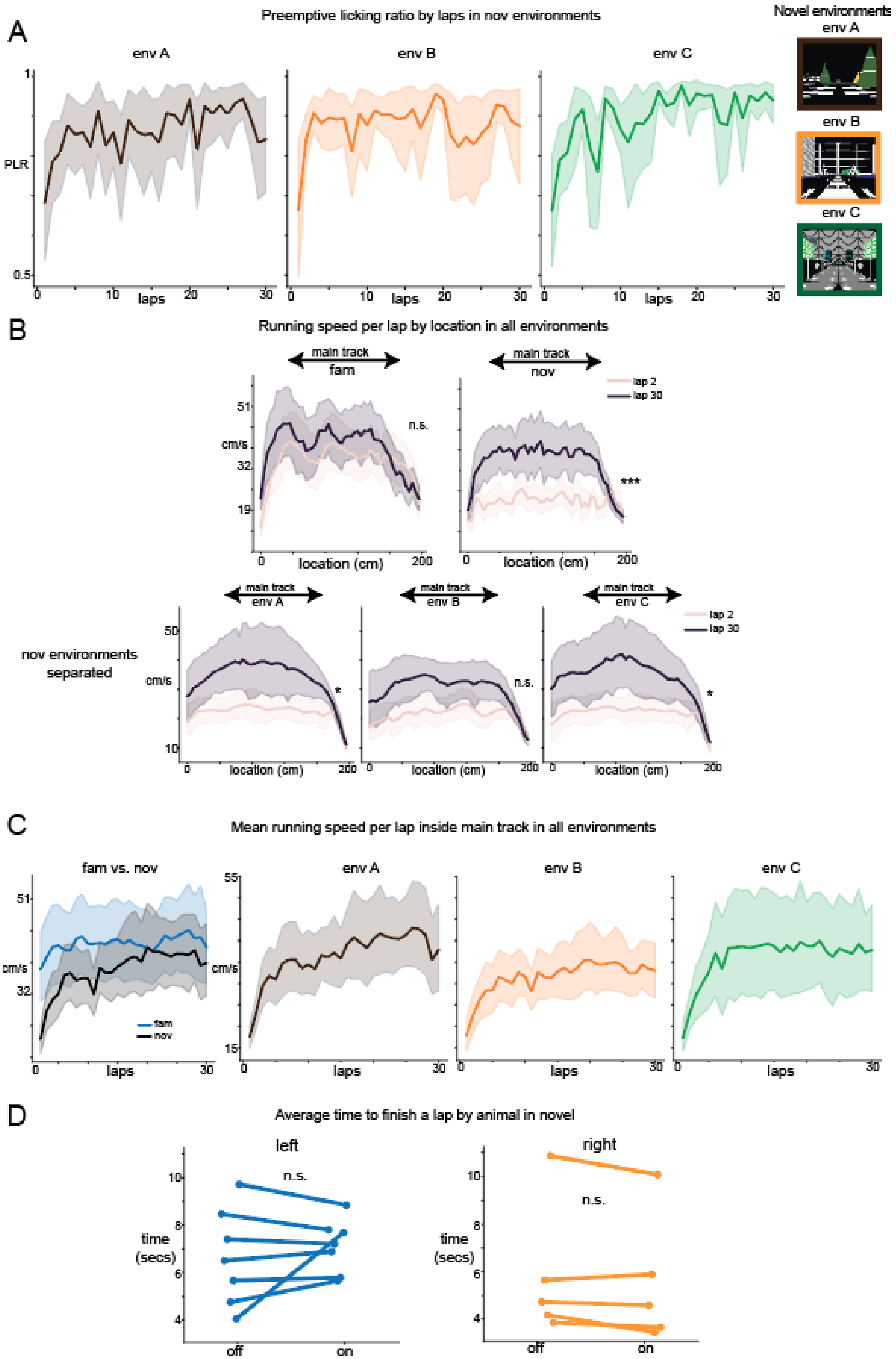


Supp Fig1. Animal behavior in familiar and novel environments.

A. Preemptive licking ratio by laps separated in 3 different novel environments averaged across all animals (N=12).
B. Top: Running speed from all animals (n=12) across all locations before rewards within lap 2 and lap 30 in familiar and 3 novel environments combined. Main track region here included track locations after consummatory licks and before pre-reward zone. Average velocity in lap 2 and lap 30 was not significantly different in familiar environment (Wilcoxon test $p=0.23$ after Bonferroni correction, $n=12$), but significantly different in novel environment (Wilcoxon test $p<0.001$ after Bonferroni correction, $n=12$). Bottom: Novel environments separated. Wilcoxon test p in three environments p=0.03, 0.06, 0.01 after Bonferroni correction, $n=12$.
C. Running speed averaged across main track region over each lap in all environments.
D. Average time elapsed (in seconds) for running one lap under optogenetic inhibition and before turning on opto across all mice. CA3 Optogenetic inhibition did not alter the mean running speed of animals in novel environments. Frames recorded during and after reward zone were not included. Wilcoxon test between opto off and on, left: $p=0.69$, $n=7$; Right: $p=0.31$, $n=5$. Between off and on pooled between all animals: Wilcoxon test $p=0.79$, $n=12$.

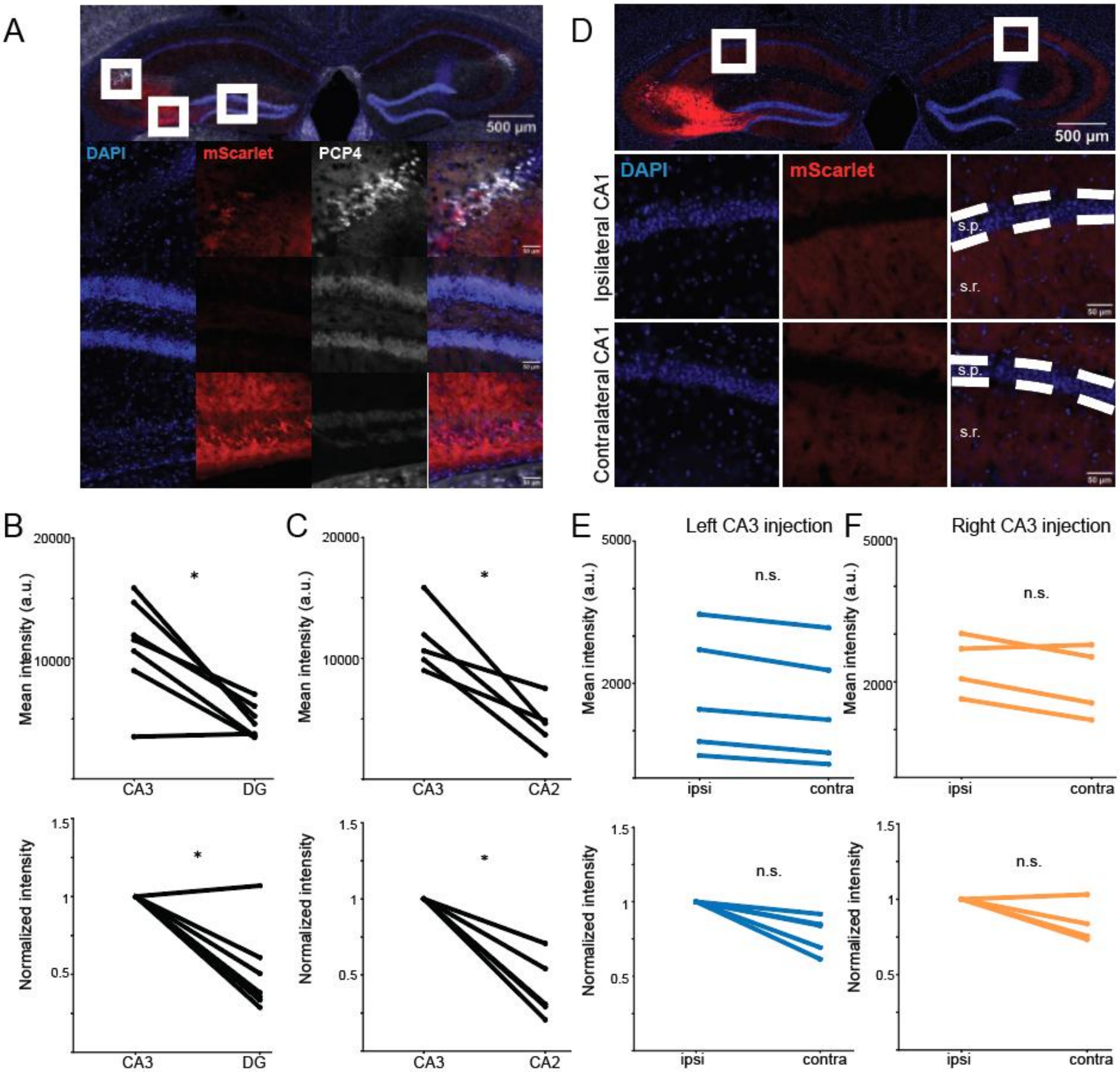


Supp Fig2. There is limited off-target eOPN3 expression in DG granule cells and CA2 pyramidal cells compared to CA3 somas.

A. (Top) Representative immunofluorescent image of the hippocampus from a Grik4-Cre mouse injected with eOPN3-mScarlet into the left CA3. Magnified view of CA2 (middle row) and DG (bottom row) from the left and right white square in the top image, respectively. Shows labeling of cell nuclei with DAPI (blue), the opsin fluorescent tag mScarlet (red), and CA2 pyramidal cells with PCP4 (white).

B. (Top) Mean intensity of mScarlet within CA3 pyramidal cells and DG granule cells within the injected hemisphere. Each dot represents the mean intensity calculated from three 30 µm tissue sections obtained from a single mouse (n = 7). (Bottom) Normalized intensity of mScarlet within DG granule cells. Mean values are normalized to the intensity in CA3 pyramidal cells for each mouse (n = 7). One-sided Wilcoxon signed-rank test (stat = 27.0 p = 0.016)

C. (Top) Mean intensity of mScarlet within CA3 and CA2 pyramidal cells within the injected

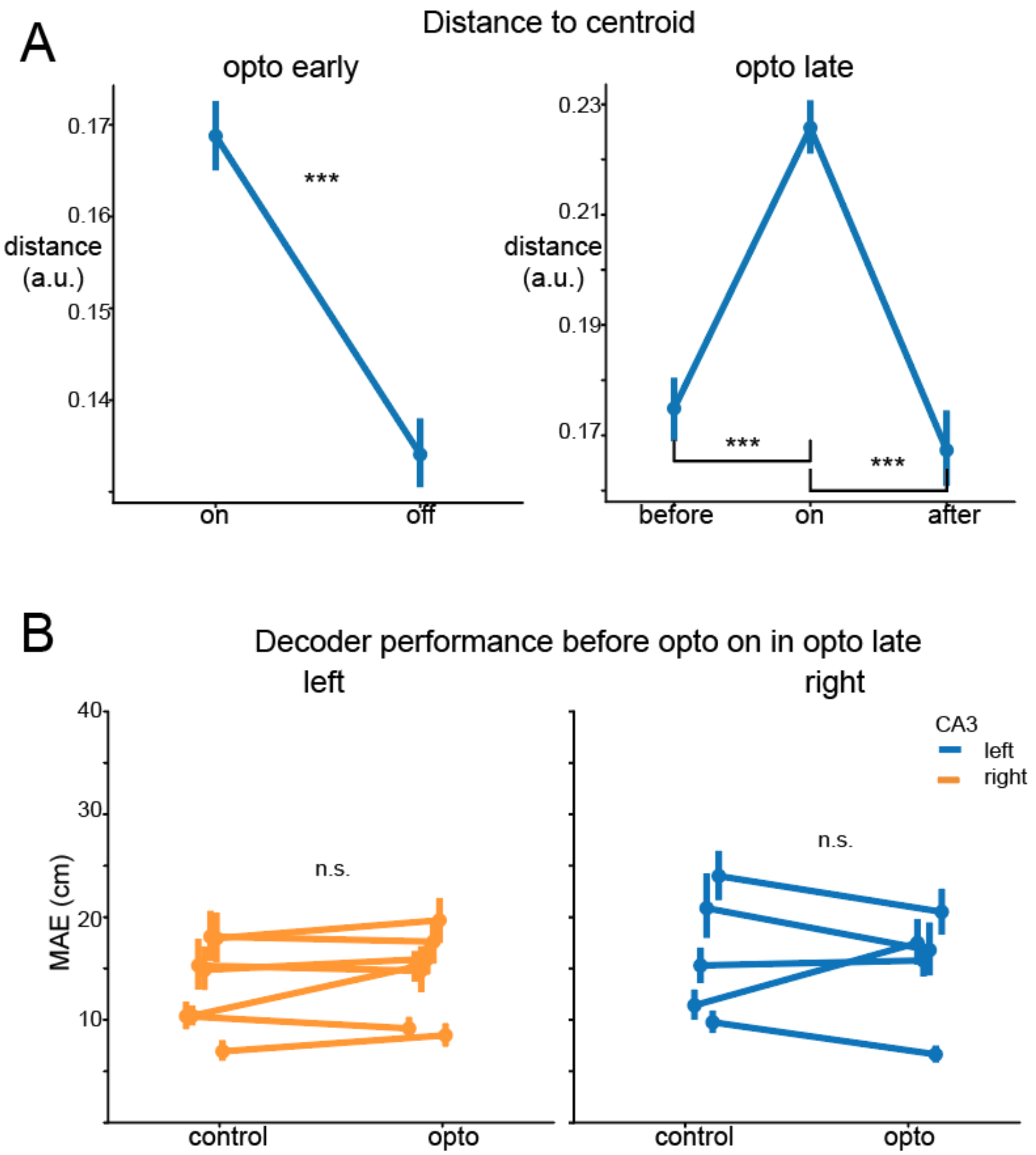


Supp Fig.3. Right optogenetic inhibition significantly increased the distance to centroid in the same example animal as Fig4, representing a less stereotypical neural activity to location on the track under opto.

A. Distance to centroid in opto early and opto late. Same example animal as (Fig4. a). See methods for calculating distance to centroid. Left: Optogenetic inhibition significantly increased distance to centroid than off in opto early. (Mann whitney u stats= 152935071, $p<0.001$). Right: Turning on optogenetic inhibition significantly increased distance to centroid than before and after. (Before vs. on: $p<0.001$; after vs. on: $p<0.001$).

B. Decoder performance before turning on optogenetic inhibition in opto late and control condition. No significant difference in decoder performance between opto and control were found in both left and right CA3 (Wilcoxon signed rank test comparing within animal difference after Bonferroni correction (*2) left: $p=0.16$, n observations=1470; right: $p=0.5$, n observations=1000). Decoder performance trained using the same protocol (see methods for details) tested in 5 laps before opto on in opto late condition.

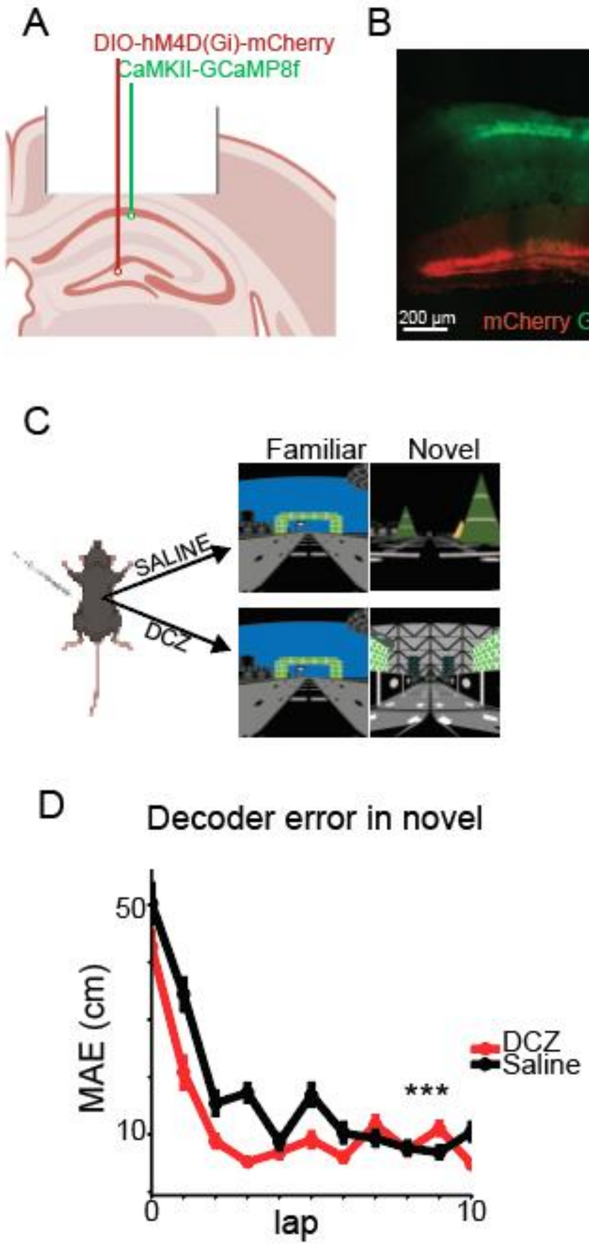


Supp Fig4. Effects of optogenetic inhibition were not driven by off-target labeling of dentate gyrus granule cells.

A-C) protocol for chemogenetic inhibition of granule cells and CA1 imaging in the right hemisphere (adapted from GoodSmith et al., 2025)[57].

A) A cre-dependent inhibitory DREADD virus was injected into the DG, and GCaMP8f was injected into dorsal right CA1.

B) mCherry expression can be seen selectively in the granule cell layer and mossy fiber projections to CA1, while GCaMP expression is restricted to CA1.

C) On two seperate days, mice (n=4) were injected with DCZ (to inhibit granule cells) or saline (as a within-animal control). On each day, mice were exposed to a novel and familiar environment in ~5.5 minute blocks.
D) Effect of granule cell inhibition on decoding error (similar to Fig 2). In contrast to the increased decoding error observed following optogenetic inhibition of CA3 activity in the early phase, granule cell inhibition significantly decreased decoding error compared to control. (n observations = 1694, ranksum test p <0.001)

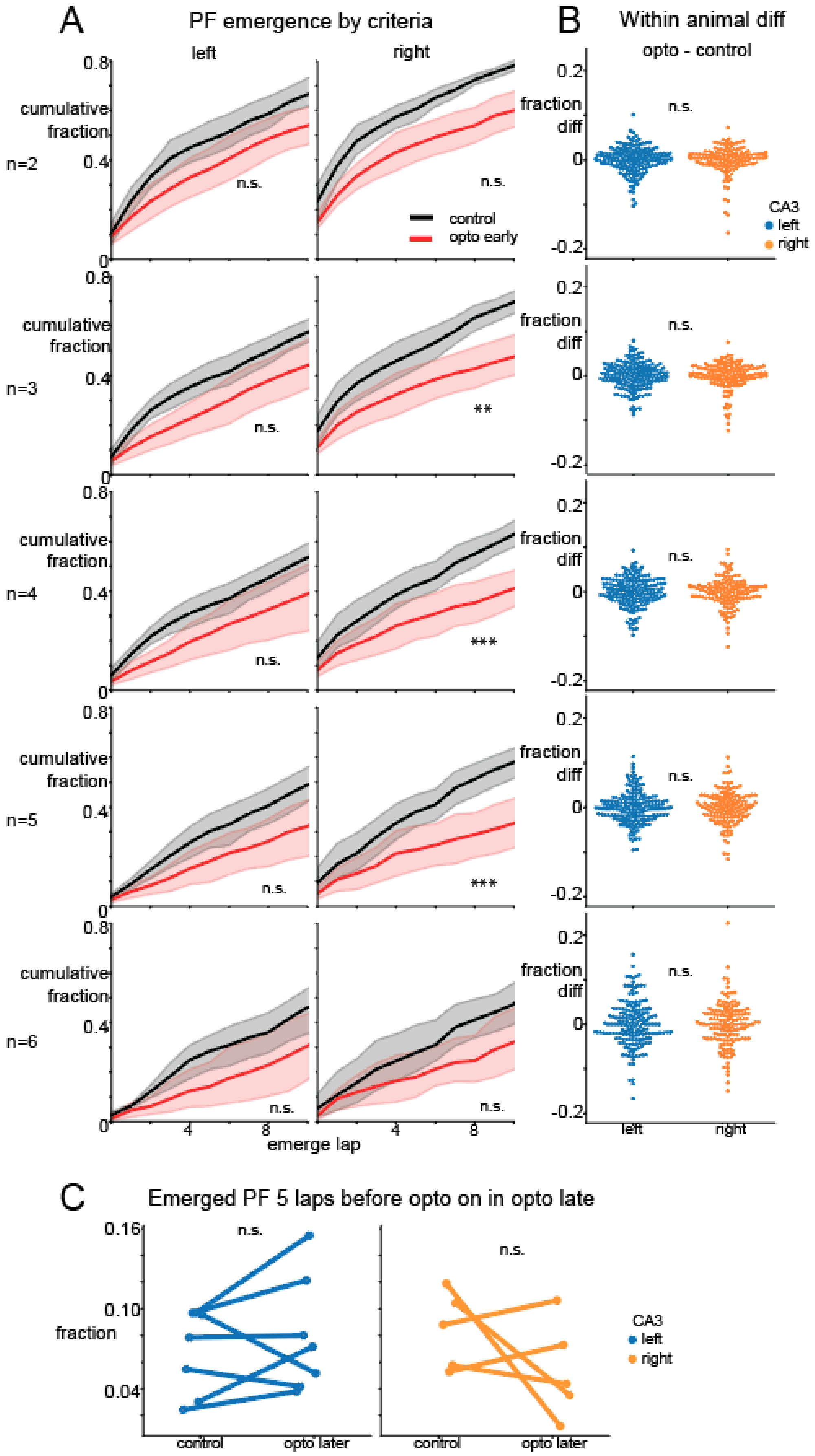

A
PF emergence by criteria
left
right
B
Within animal diff
opto - control
cumulative fraction
fraction diff
n=2
n=3
n=4
n=5
n=6
0.8
0.4
0
0.2
-0.2
n.s.
**
***
control
opto early
CA3
left
right
4
8
emerge lap
C
Emerged PF 5 laps before opto on in opto late
fraction
0.16
0.10
0.04
control
opto later

Supp Fig5. Left and right CA3 input inhibition similarly delayed the emergence of CA1 place cells, regardless of emergence criteria.

A. Cumulative histogram of fractions of place fields emerged by lap number comparing opto early (red) with control (black) for first 10 laps under optogenetic inhibition by difference emergence criteria. Emergence lap was defined as the first lap in N out of 6 consecutive laps of within-field firing. P values reported were after Bonferroni correction (*10). N=2, left: Wilcoxon signed-rank stats=701, p =1.2; right: stats= 361.5, p=0.07. N=3, left: stats=612, p=0.26; right: stats=262, p<0.01; N=4, left: stats=588, p=0.16; right: stats=209, p<0.001; N=5, left: stats=543, p=0.062; right: stats=190, p<0.001; N=6, left: stats=506, p=0.11; right: stats=371, p=0.26.
B. Within-animal emergence fraction difference between control and opto. Each dot represented one lap: below 0 meant the lap had less fraction of PF emerged under opto than that under lap-matched control, which reflected that optogenetic inhibition delayed the emergence of place fields in that lap. No significant difference found between left and right under any criteria of N. P values reported were after Bonferroni correction (*5). N=2, Mann whitney U stats=1653, p=1.40; N=3, Mann whitney U stats=1702, p=1.23; N=4, T stats = 1.29, p=1; N=5, T stats=1.74, p=0.41; N=6, T stats=0.24, p=4.05.
C. Fractions of place fields emerged 5 laps before turning on optogenetic in opto late by animal. No significant differences were found between control and opto in the left or right group. (linear mixed effects model, no significant effects, CA3 p=0.99, opto p=0.46, CA3 x opto p=0.087. Wilcoxon signed-rank test left: p=0.375, n=7, right: p=0.625, n=5.)

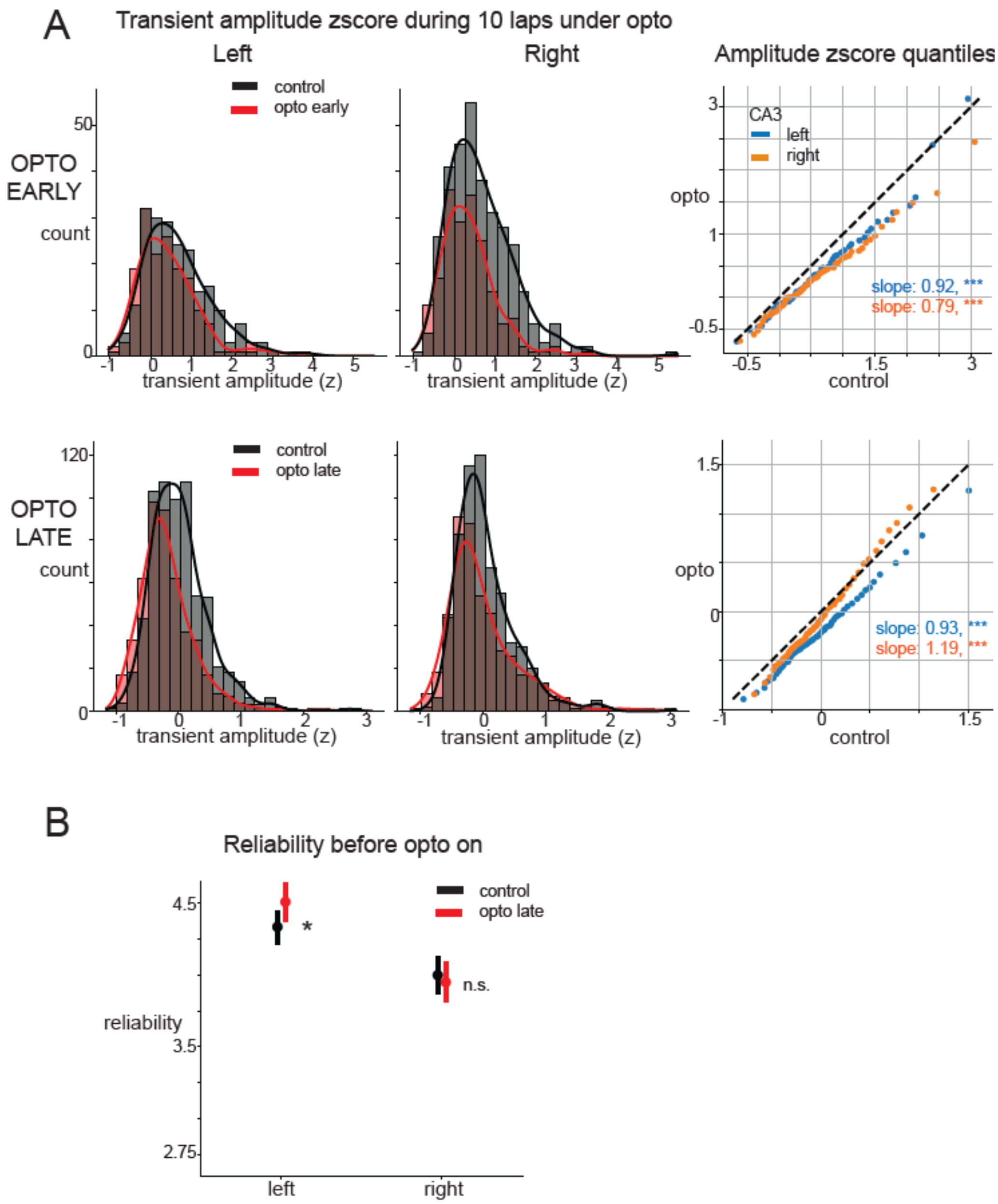


Supp Fig6. Characterizing place cell amplitude and reliability metrics.

A. In-field transient amplitude distribution under opto and control in histogram and qqplot. Top: In opto early, inhibiting left and right CA3 axons during the early phase disproportionately reduced high amplitude transients. Low amplitude transients were not reduced. Left n transients opto=166, control=211; right n transients opto=188, control=332. Quantile-quantile plot compared the distribution of z-scored transient amplitudes in control and opto. A linear regression was fitted to the qqplot. Left: R=0.98, $p<0.001$, slope=0.92, intercept=-0.17; Right: R=0.99, $p<0.001$, slope=0.79, intercept=-0.17. Dashed line showed if opto and control had the same distribution for reference, slope smaller than 1 reflects a bigger effect on higher amplitude transients, 1 reflects same effect on all transients regardless of amplitude, bigger than 1 reflects a bigger effect on lower amplitude transients. Bottom: Same as top in opto late. In the later phase, left CA3 input inhibition equally reduced all transients independent of their amplitude. However, right CA3 input inhibition mostly affected low amplitude transients. Left n transients opto=468, control=635; right n transients opto=465, control=603; qqplot. Left: R=0.99, p<0.001, slope=0.93, intercept=-0.2; Right: R=0.99, p<0.001, slope=1.19, intercept=-0.05.

B. In-field reliability 5 laps before opto on in opto late condition as control. Right CA3 animals showed no difference from controls. left CA3 animals exhibited slightly higher reliability than controls - opposite to the direction caused by inhibition. Left: N place fields control=781, opto=612, Mann whitney U p=0.034 after Bonferroni correction (*2); Right: N place fields control=818, opto=656, p=0.88.

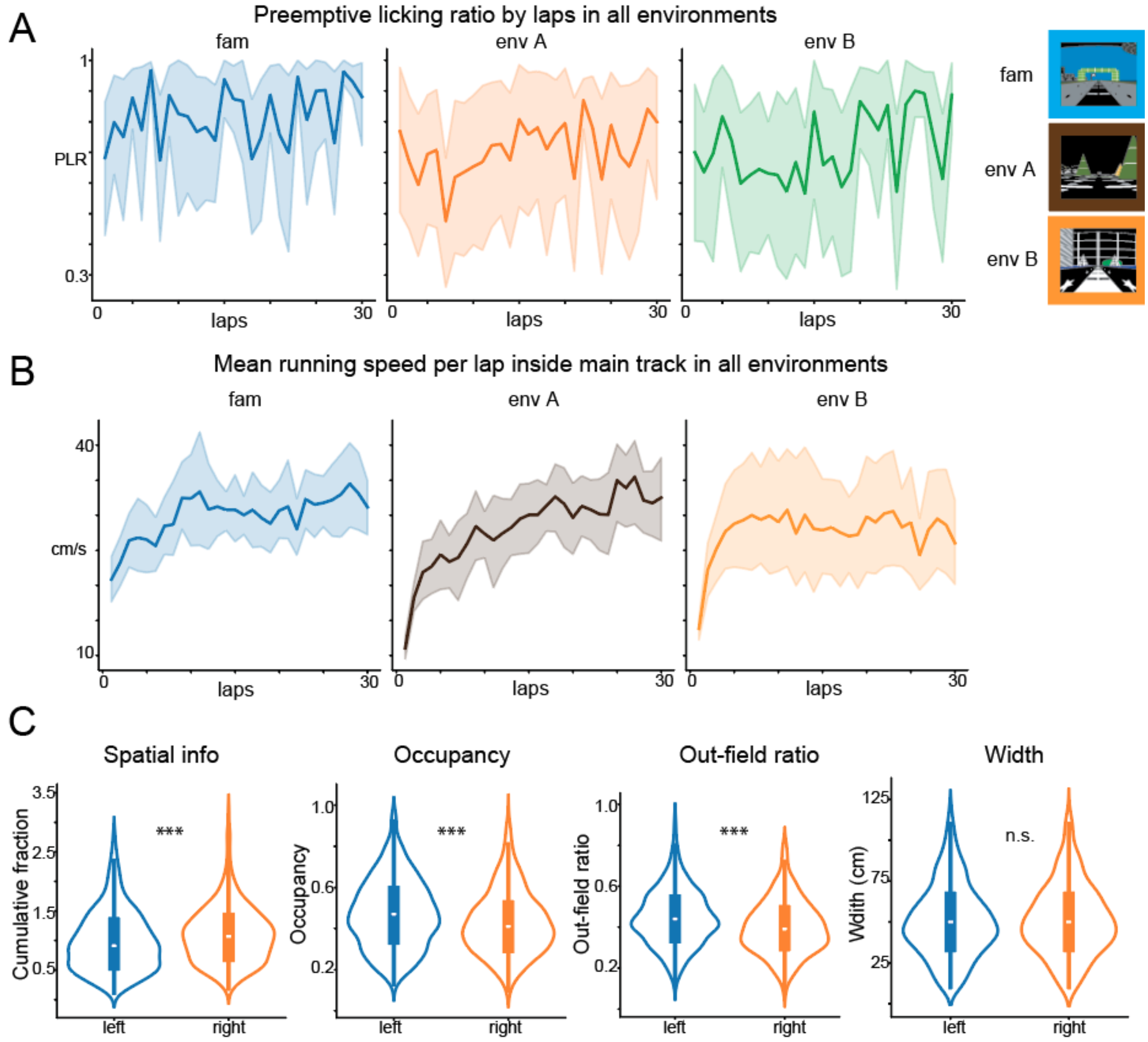


Supp Fig7. Axon animal behavior and place fields features.

A. Mean preemptive licking ratio calculated the same way as Fig1 in axon animals (n=8) in all environments.

B. Mean running speed inside the same main track region as Supp. Fig.1 in axon animals (n=8) over laps in all environments.

C. Left and right CA3 axon place fields (left n=813, right n=530) feature comparison. Right CA3 place fields contained significantly more spatial information. Mann Whitney U stats=450693, p<0.001. Left CA3 axon place fields had significantly longer occupancy than right CA3 axon place fields. Mann Whitney U stats=607665, p<0.001. Left CA3 axon place fields had significantly more out-field firing ratio than right CA3 axon place fields. Mann Whitney U stats=633514, p<0.001. Left and right CA3 axon place fields were not significantly different from each other. Mann Whitney U stats=514315, p=0.38

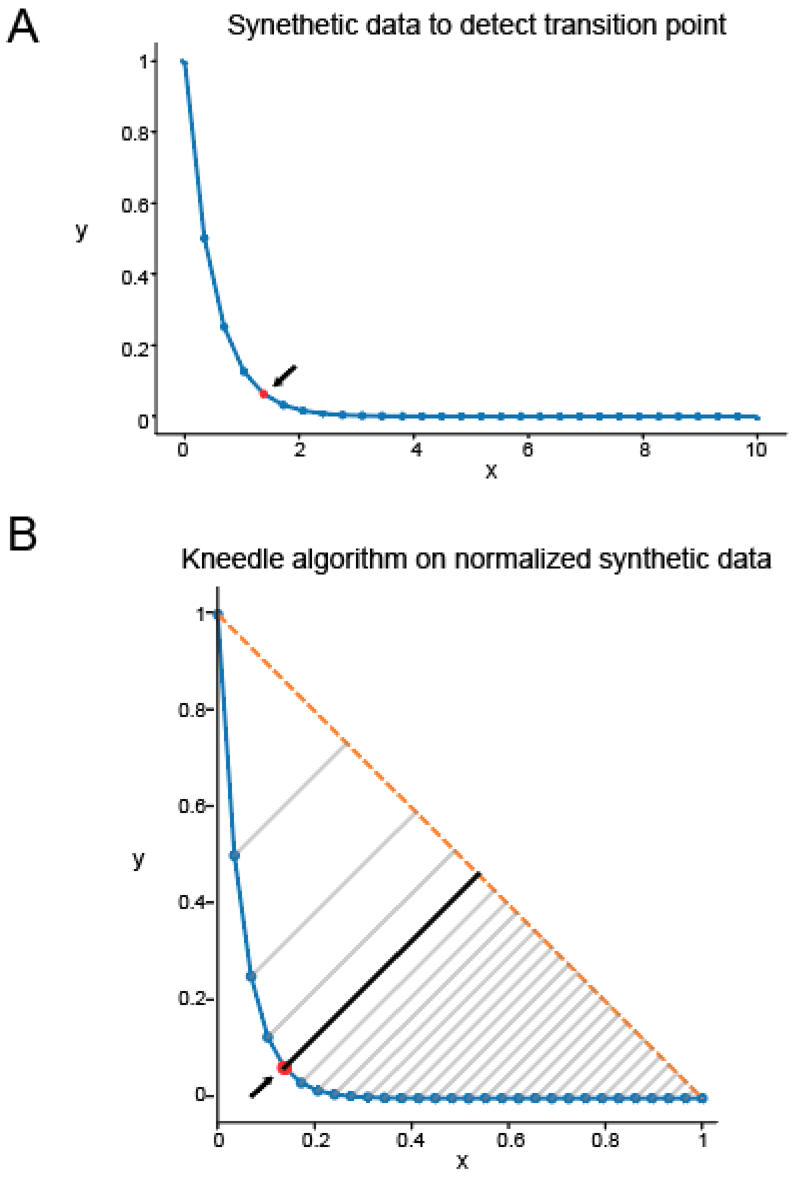

Supp Fig8 Illustration of kneedle algorithm.

A. Example synthetic data with an exponential decay profile used to illustrate detection of a transition point. Arrow pointing at the transition point (red).
B. Same dataset after normalization and application of the Kneedle algorithm. The method identifies the point of maximum deviation between the data curve (blue) and the diagonal line (orange dashed) connecting the start and end of the curve (smoothing as needed when using real data). The knee point corresponds to the location with the largest perpendicular distance (black line) from this diagonal, highlighted by the red point and arrow.

**Funding.** This work was supported by The Whitehall Foundation, The Searle Scholars Program, The Sloan Foundation, The University of Chicago Institute for Neuroscience start-up funds and the National Institute for Health (1DP2NS111657-01, 1RF1NS127123-01, 1R21NS128822-01, 1R01MH136274-01) awarded to M.S.

**Author contributions.** A.J. and M.S. conceived and designed the experiments. A.J. collected the optogenetic and axon data. A.J. analyzed the data. J.R.M. and A.T. collected the histology and immunofluorescence data. J.R.M. analyzed the histology and immunofluorescence data. D.G. collected and analyzed the dentate gyrus data. A.J. and M.S. interpreted the data and wrote the original manuscript. A.J., D.G., J.R.M., A.T., M.S. reviewed and edited the manuscript. M.S. supervised the research and obtained funding.

**Declaration of interests.** The authors declare no competing interests.

**Data and code availability**. The original codes have been deposited at https://github.com/anqijiang/opto_analysis and will be publicly available on the date of publication. Raw imaging data is large and not feasible for upload to an online repository but is available upon request to the lead contact at. Processed source data for all figures and associated statistical analysis will be uploaded to Box and provided in the final version of the paper. Any additional information required to reanalyze the data reported in this paper is available from the lead contact upon request.

### References

1. Eichenbaum, H., Dudchenko, P., Wood, E., Shapiro, M., and Tanila, H. (1999). The Hippocampus, Memory, and Place Cells Is It Spatial Memory or a Memory Space? Neuron *23*, 209–*226*. https://doi.org/*10*.10*16*/s08*96*-6273(00)80773-4.
2. Robinson, N.T.M., Descamps, L.A.L., Russell, L.E., Buchholz, M.O., Bicknell, B.A., Antonov, G.K., Lau, J.Y.N., Nutbrown, R., Schmidt-Hieber, C., and Häusser, M. (2020). Targeted Activation of Hippocampal Place Cells Drives Memory-Guided Spatial Behavior. Cell *183*, 1586-1599.e10. https://doi.org/10.1016/j.cell.2020.09.061.
3. O'Keefe, J., and Nadel, L. (1979). Précis of O'Keefe & Nadel's The hippocampus as a cognitive map. Behav. Brain Sci. 2, 487–494. https://doi.org/10.10*17*/s0*14*0525x00063949.
4. Leussis, M.P., and Bolivar, V.J. (2006). Habituation in rodents: A review of behavior, neurobiology, and genetics. Neurosci. Biobehav. Rev. *30*, 1045–1064. https://doi.org/10.1016/j.neubiorev.2006.03.006.
5. Wilkinson, J.L., Herrman, L., Palmatier, M.I., and Bevins, R.A. (2006). Rats' novel object interaction as a measure of environmental familiarity. Learn. Motiv. *37*, *13*1–148. https://doi.org/10.1016/j.lmot.2005.04.001.
6. Bevins, R.A., Koznarova, J., and Armiger, T.J. (2001). Environmental Familiarization in Rats: Differential Effects of Acute and Chronic Nicotine. Neurobiol. Learn. Mem. *75*, 63–76. https://doi.org/10.1006/nlme.1999.3955.
7. Dong, C., Madar, A.D., and Sheffield, M.E.J. (2021). Distinct place cell dynamics in CA1 and CA3 encode experience in new environments. Nat. Commun. *12*, *297*7. https://doi.org/10.1038/s*41*467-021-23260-3.
8. Berners-Lee, A., Feng, T., Silva, D., Wu, X., Ambrose, E.R., Pfeiffer, B.E., and Foster, D.J. (2022). Hippocampal replays appear after a single experience and incorporate greater detail with more experience. Neuron *110*, 1829-1842.e5. https://doi.org/10.1016/j.neuron.2022.03.010.
9. Cohen, J.D., Bolstad, M., and Lee, A.K. (2017). Experience-dependent shaping of hippocampal CA1 intracellular activity in novel and familiar environments. eLife 6, e23040. https://doi.org/10.7554/elife.23040.
10. Chiu, Y., Dong, C., Krishnan, S., and Sheffield, M.E.J. (2023). The Precision of Place Fields Governs Their Fate across Epochs of Experience. eNeuro 10, ENEURO.0261-23.2023. https://doi.org/10.1523/eneuro.0261-23.2023.
11. Zemla, R., Moore, J.J., Hopkins, M.D., and Basu, J. (2022). Task-selective place cells show behaviorally driven dynamics during learning and stability during memory recall. Cell Rep. 41, *111*700. https://doi.org/10.1016/j.celrep.2022.111700.
12. Sheffield, M.E.J., Adoff, M.D., and Dombeck, D.A. (2017). Increased Prevalence of Calcium Transients across the Dendritic Arbor during Place Field Formation. Neuron 96, 490-*50*4.e5. https://doi.org/10.1016/j.neuron.2017.09.029.
13. Geiller, T., Sadeh, S., Rolotti, S.V., Blockus, H., Vancura, B., Negrean, A., Murray, A.J., Rózsa, B., Polleux, F., Clopath, C., et al. (2022). Local circuit amplification of spatial selectivity in the hippocampus. Nature *601*, *105–109*. https://doi.org/10.1038/s41586-021-04169-9.
14. Bittner, K.C., Milstein, A.D., Grienberger, C., Romani, S., and Magee, J.C. (2017). Behavioral time scale synaptic plasticity underlies CA1 place fields. Science *357*, 1033–1036. https://doi.org/10.1126/science.aan3846.

15. Bittner, K.C., Grienberger, C., Vaidya, S.P., Milstein, A.D., Macklin, J.J., Suh, J., Tonegawa, S., and Magee, J.C. (2015). Conjunctive input processing drives feature selectivity in hippocampal CA1 neurons. Nat. Neurosci. 18, 1133–1142. https://doi.org/10.1038/nn.4062.
16. Click or tap here to enter text.
Madar, A.D., Jiang, A., Dong, C., and Sheffield, M.E.J. (2025). Synaptic plasticity rules driving representational shifting in the hippocampus. Nat. Neurosci., 1–13. https://doi.org/10.1038/s41593-025-01894-6.55.
17. Priestley, J.B., Bowler, J.C., Rolotti, S.V., Fusi, S., and Losonczy, A. (2022). Signatures of rapid plasticity in hippocampal CA1 representations during novel experiences. Neuron 110, 19*78*-1992.e6. https://doi.org/10.1016/j.neuron.2022.03.026.
18. Kesner, R.P. (2007). Behavioral functions of the CA3 subregion of the hippocampus. Learn. Mem. 14, 771–781. https://doi.org/10.1101/lm.688207.
19. Nakazawa, K., Quirk, M.C., Chitwood, R.A., Watanabe, M., Yeckel, M.F., Sun, L.D., Kato, A., Carr, C.A., Johnston, D., Wilson, M.A., et al. (2002). Requirement for Hippocampal CA3 NMDA Receptors in Associative Memory Recall. Science 297, 211–218. https://doi.org/10.1126/science.1071795.
20. Li, X. - G., Somogyi, P., Ylinen, A., and Buzsáki, G. (1994). The hippocampal CA3 network: An in vivo intracellular labeling study. J. Comp. Neurol. *339*, 181–208. https://doi.org/10.1002/cne.903390204.
21. Kohl, M.M., Shipton, O.A., Deacon, R.M., Rawlins, J.N.P., Deisseroth, K., and Paulsen, O. (2011). Hemisphere-specific optogenetic stimulation reveals left-right asymmetry of hippocampal plasticity. Nat. Neurosci. 14, 1413–1415. https://doi.org/10.1038/nn.2915.
22. Shinohara, Y., Hirase, H., Watanabe, M., Itakura, M., Takahashi, M., and Shigemoto, R. (2008). Left-right asymmetry of the hippocampal synapses with differential subunit allocation of glutamate receptors. Proc. Natl. Acad. Sci. 105, 19498–19503. https://doi.org/10.1073/pnas.0807461105.
23. Shipton, O.A., El-Gaby, M., Apergis-Schoute, J., Deisseroth, K., Bannerman, D.M., Paulsen, O., and Kohl, M.M. (2014). Left–right dissociation of hippocampal memory processes in mice. Proc. Natl. Acad. Sci. 111, 15238–15*243*. https://doi.org/10.1073/pnas.1405648111.
24. Guan, H., Middleton, S.J., Inoue, T., and McHugh, T.J. (2021). Lateralization of CA1 assemblies in the absence of CA3 input. Nat. Commun. 12, 6114. https://doi.org/10.1038/s41467-021-26389-3.
25. El-Gaby, M., Zhang, Y., Wolf, K., Schwiening, C.J., Paulsen, O., and Shipton, O.A. (2016). Archaerhodopsin Selectively and Reversibly Silences Synaptic Transmission through Altered pH. Cell Rep. 16, 2259–2268. https://doi.org/10.1016/j.celrep.2016.07.057.
26. Song, D., Wang, D., Yang, Q., Yan, T., Wang, Z., Yan, Y., Zhao, J., Xie, Z., Liu, Y., Ke, Z., et al. (2020). The lateralization of left hippocampal CA3 during the retrieval of spatial working memory. Nat. Commun. 11, 2901. https://doi.org/10.1038/s41467-020-16698-4.
27. El-Gaby, M., Reeve, H.M., Lopes-dos-Santos, V., Campo-Urriza, N., Perestenko, P.V., Morley, A., Strickland, L.A.M., Lukács, I.P., Paulsen, O., and Dupret, D. (2021). An emergent neural coactivity code for dynamic memory. Nat. Neurosci. 24, 694–704. https://doi.org/10.1038/s41593-021-00820-w.
*28*. Kawakami, R., Shinohara, Y., Kato, Y., Sugiyama, H., Shigemoto, R., and Ito, I. (2003). Asymmetrical Allocation of NMDA Receptor ε2 Subunits in Hippocampal Circuitry. Science *300*,

990–994. https://doi.org/10.1126/science.1082609.
29. Jordan, J.T. (2020). The rodent hippocampus as a bilateral structure: A review of hemispheric lateralization. Hippocampus 30, 278–292. https://doi.org/10.1002/hipo.23188.
30. Mahn, M., Saraf-Sinik, I., Patil, P., Pulin, M., Bitton, E., Karalis, N., Bruentgens, F., Palgi, S., Gat, A., Dine, J., et al. (2021). Efficient optogenetic silencing of neurotransmitter release with a mosquito rhodopsin. Neuron 109, 1621-1635.e8. https://doi.org/10.1016/j.neuron.2021.03.013.
31. Satopää, V., Albrecht, J., Irwin, D., and Raghavan, B. (2011). Finding a “Kneedle” in a Haystack: Detecting Knee Points in System Behavior. 2011 31st Int. Conf. Distrib. Comput. Syst. Work., 166–171. https://doi.org/10.1109/icdcsw.2011.20.
32. Davoudi, H., and Foster, D.J. (2019). Acute silencing of hippocampal CA3 reveals a dominant role in place field responses. Nat. Neurosci. 22, 337–342. https://doi.org/10.1038/s41593-018-0321-z.
33. Zutshi, I., Valero, M., Fernández-Ruiz, A., and Buzsáki, G. (2022). Extrinsic control and intrinsic computation in the hippocampal CA1 circuit. Neuron 110, 658-673.e5. https://doi.org/10.1016/j.neuron.2021.11.015.
34. Guo, W., Zhang, J.J., Newman, J.P., and Wilson, M.A. (2024). Latent learning drives sleep-dependent plasticity in distinct CA1 subpopulations. Cell Rep. 43, 115028. https://doi.org/10.1016/j.celrep.2024.115028.
35. Yang, W., Sun, C., Huszár, R., Hainmueller, T., Kiselev, K., and Buzsáki, G. (2024). Selection of experience for memory by hippocampal sharp wave ripples. Science *383*, 1478–1483. https://doi.org/10.1126/science.adk8261.
36. Nieh, E.H., Schottdorf, M., Freeman, N.W., Low, R.J., Lewallen, S., Koay, S.A., Pinto, L., Gauthier, J.L., Brody, C.D., and Tank, D.W. (2021). Geometry of abstract learned knowledge in the hippocampus. Nature *595*, 80–84. https://doi.org/10.1038/s41586-021-03652-7.
37. Jones, E.A.A., and Giocomo, L.M. (2023). Neural ensembles in navigation: From single cells to population codes. Curr. Opin. Neurobiol. 78, 102665. https://doi.org/10.1016/j.conb.2022.102665.
38. Schneider, S., Lee, J.H., and Mathis, M.W. (2023). Learnable latent embeddings for joint behavioural and neural analysis. Nature *617*, 360–368. https://doi.org/10.1038/s41586-023-06031-6.
39. McKenzie, S., Huszár, R., English, D.F., Kim, K., Christensen, F., Yoon, E., and Buzsáki, G. (2021). Preexisting hippocampal network dynamics constrain optogenetically induced place fields. Neuron 109, 1040-1054.e7. https://doi.org/10.1016/j.neuron.2021.01.011.
40. Keinath, A.T., Nieto-Posadas, A., Robinson, J.C., and Brandon, M.P. (2020). DG–CA3 circuitry mediates hippocampal representations of latent information. Nat. Commun. 11, 3026. https://doi.org/10.1038/s41467-020-16825-1.
41. Ji, D., and Wilson, M.A. (2008). Firing Rate Dynamics in the Hippocampus Induced by Trajectory Learning. J. Neurosci. 28, 4679–4689. https://doi.org/10.1523/jneurosci.4597-07.2008.
42. Kitamura, T., Ogawa, S.K., Roy, D.S., Okuyama, T., Morrissey, M.D., Smith, L.M., Redondo, R.L., and Tonegawa, S. (2017). Engrams and circuits crucial for systems consolidation of a memory. Science *356*, 73–78. https://doi.org/10.1126/science.aam6808.
43. Butola, T., Hernández-Frausto, M., Blankvoort, S., Flatset, M.S., Peng, L., Hairston, A., Johnson, C.D., Elmaleh, M., Amilcar, A., Hussain, F., et al. (2025). Hippocampus shapes

entorhinal cortical output through a direct feedback circuit. Nat. Neurosci., 1–12. https://doi.org/10.1038/s41593-025-01883-9.
44. Jiang, A., Zhao, C., and Sheffield, M. (2024). A preprocessing toolbox for 2-photon subcellular calcium imaging. bioRxiv, 2024.10.04.616737. https://doi.org/10.1101/2024.10.04.616737.
45. Rebola, N., Carta, M., and Mulle, C. (2017). Operation and plasticity of hippocampal CA3 circuits: implications for memory encoding. Nat. Rev. Neurosci. 18, 208–220. https://doi.org/10.1038/nrn.2017.10.
46. Zaki, Y., and Cai, D.J. (2025). Memory engram stability and flexibility. Neuropsychopharmacology 50, 285–293. https://doi.org/10.1038/s41386-024-01979-z.
47. Geva, N., Deitch, D., Rubin, A., and Ziv, Y. (2023). Time and experience differentially affect distinct aspects of hippocampal representational drift. Neuron 111, 2357-2366.e5. https://doi.org/10.1016/j.neuron.2023.05.005.
48. Keinath, A.T., Mosser, C.-A., and Brandon, M.P. (2022). The representation of context in mouse hippocampus is preserved despite neural drift. Nat. Commun. 13, 2415. https://doi.org/10.1038/s41467-022-30198-7.
49. Khatib, D., Ratzon, A., Sellevoll, M., Barak, O., Morris, G., and Derdikman, D. (2023). Active experience, not time, determines within-day representational drift in dorsal CA1. Neuron 111, 2348-2356.e5. https://doi.org/10.1016/j.neuron.2023.05.014.
50. Krishnan, S., and Sheffield, M.E.J. (2023). Reward Expectation Reduces Representational Drift in the Hippocampus. bioRxiv, 2023.12.21.572809. https://doi.org/10.1101/2023.12.21.572809.
51. Delamare, G., Zaki, Y., Cai, D.J., and Clopath, C. (2023). Drift of neural ensembles driven by slow fluctuations of intrinsic excitability. https://doi.org/10.7554/elife.88053.2.
52. Krishnan, S., Heer, C., Cherian, C., and Sheffield, M.E.J. (2022). Reward expectation extinction restructures and degrades CA1 spatial maps through loss of a dopaminergic reward proximity signal. Nat. Commun. 13, 6662. https://doi.org/10.1038/s41467-022-34465-5.
53. Pachitariu, M., Stringer, C., Dipoppa, M., Schröder, S., Rossi, L.F., Dalgleish, H., Carandini, M., and Harris, K.D. (2017). Suite2p: beyond 10,000 neurons with standard two-photon microscopy. bioRxiv, 061507. https://doi.org/10.1101/061507.
54. Grijseels, D.M., Shaw, K., Barry, C., and Hall, C.N. (2021). Choice of method of place cell classification determines the population of cells identified. PLoS Comput. Biol. 17, e1008835. https://doi.org/10.1371/journal.pcbi.1008835.
55. Mahn, M., Saraf-Sinik, I., Patil, P., Pulin, M., Bitton, E., Karalis, N., Bruentgens, F., Palgi, S., Gat, A., Dine, J., et al. (2021). Efficient optogenetic silencing of neurotransmitter release with a mosquito rhodopsin. Neuron *109*, 1621-1635.e8. https://doi.org/10.1016/j.neuron.2021.03.013.
56. Heer, C., and Sheffield, M. (2024). Distinct catecholaminergic pathways projecting to hippocampal CA1 transmit contrasting signals during navigation in familiar and novel environments. eLife *13*, RP95213. https://doi.org/10.7554/elife.95213.56.
57. GoodSmith, D., Carson, W.H., and Sheffield, M.E.J. (2025). Complementary regulation of memory flexibility and stabilization by dentate gyrus granule cells and mossy cells. BioRxiv. https://doi.org/10.1101/2025.09.05.674488.
